# Identification of Malignant Peripheral Nerve Sheath Tumor subtypes with distinct genomic identities

**DOI:** 10.64898/2026.03.31.715523

**Authors:** Miriam Magallón-Lorenz, Juana Fernández-Rodríguez, Helena Mazuelas, Itziar Uriarte-Arrazola, Sara Ortega-Bertrán, Edgar Creus-Bachiller, Judit Farrés-Casas, Aleix Mendez, Eva Rodríguez, Mariona Suñol, Carlota Rovira, Raquel Arnau, Tulio Silva, Juan Carlos Lopez-Gutierrez, Alicia Castañeda, Isabel Granada, Alba Hernández-Gallego, Gustavo Tapia, Maria Saigí, Marc Cucurull, Ignacio Blanco, Claudia Valverde, Cleofé Romagosa, Héctor Salvador, Conxi Lázaro, Meritxell Carrió, Eduard Serra, Bernat Gel

## Abstract

Malignant peripheral nerve sheath tumors (MPNSTs) are aggressive soft-tissue sarcomas arising sporadically or in people with neurofibromatosis type 1 (NF1). Their marked heterogeneity challenges diagnosis and has hampered an integrative view of MPNST molecular pathogenesis. Here, a thorough whole-genome and transcriptome analysis of MPNSTs and the re-analysis of a large independent cohort allowed us to identify three molecular subtypes of MPNSTs (G1-G3) with distinct genomic identities and clinicopathological features. Furthermore, it provided a simple and unifying model of MPNST development, defining a distinct progression path for each group. This work uncovers new genomic aspects of MPNSTs, including the identification of recurrent copy-neutral loss of heterozygosity regions, distinct copy-number profiles among G1-G3, and *CDKN2A*-inactivating translocations in pre-malignant lesions (ANNUBPs). Altogether, these analyses overcome the dominant influence of PRC2 status in MPNST classification and provide a framework for their differential diagnosis and potential precision oncology treatment.

**SIGNIFICANCE:** MPNST is a highly heterogeneous soft-tissue sarcoma with difficult clinical management and no effective systemic therapies. This work defines three molecular subtypes of MPNSTs with distinct development paths and histological and clinical characteristics with potential impact on translational studies and subtype-tailored treatments.

## INTRODUCTION

Malignant Peripheral Nerve Sheath Tumors (MPNSTs) are aggressive soft-tissue sarcomas that may arise sporadically or, at a much higher risk, in patients with Neurofibromatosis Type 1 (NF1) (1–3) and are the leading cause of death in these patients. (1,4). The prognosis remains discouraging, with high rates of local recurrence and metastatic spread (5–7). The diagnosis of MPNST might be challenging due to the existence of heterogeneity in the histological presentation and biological features, and the existence of other tumor entities that share overlapping histological characteristics (8–11), impacting the clinical management (12–14). Complete tumor resection with wide margins is essential for a good prognosis, like for many other soft tissue sarcomas, often followed by radiation and/or chemotherapy (15).

Genomically, MPNSTs have hyperploid and highly rearranged genomes with a low mutation burden (16–18). In the context of NF1, it is well established that MPNST development is first driven by the inactivation of tumor suppressor genes (TSGs) along tumor progression. A pre-existing benign plexiform neurofibroma (pNF) is characterized by the complete loss of *NF1* (19–21). Then, *CDKN2A* inactivation occurs in an intermediate pre-malignant discrete nodular lesion with uncertain biological capacity termed atypical neurofibroma or ANNUBP (10,21–24). In fact, *CDKN2A* has been found recurrently inactivated by translocations in a hotspot region close to exon 2 in MPNST, pointing out a path for malignant progression (25). Additionally, MPNSTs lose the function of the Polycomb Repressive complex 2 (PRC2), by inactivation of *SUZ12* or *EED* (26–29).

Previous large-scale studies on genomic, transcriptomic and/or epigenomic analyses provided a highly valuable molecular characterization of MPNSTs, trying to better understand existing MPNST heterogeneity and MPNST formation (18,30–33). However, despite these significant efforts, there is still little integration of findings among different works and a comprehensive model of MPNST genesis that covers MPNST heterogeneity is still missing.

In this study, we performed a fine and dedicated deep whole-genomic and transcriptomic analysis of a discovery cohort of 20 diagnosed MPNSTs and 7 pre-malignant ANNUBPs. This analysis allowed the identification of several genomic characteristics of MPNSTs, such as the existence of recurrent copy-neutral loss of heterozygosity (LOH) regions, *CDKN2A*-inactivating translocations already in ANNUBPs and highly recurrent copy number alterations. After an additional in-depth de novo re-analysis of 50 additional samples from the Genomics of MPNST (GeM) consortium dataset (18) we applied the gained genomic understanding to the whole set of MPNSTs. Altogether, we identified three robust MPNST genomic groups that associated with clinicopathological characteristics and built a simple three-step model (initiation, progression, and stabilization) of MPNST genesis. These findings provide a new framework for MPNST stratification, moving beyond the dominant effect of PRC2 status, linking fundamental genomic architecture to clinical behavior and therapeutic vulnerability.

## RESULTS

### Transcription factor expression uncovers heterogeneity in diagnosed MPNSTs, revealing two main clusters

Accurate classification of tumors diagnosed as MPNST remains challenging, in part because multiple entities can mimic MPNST histology. To assess whether integrated genomics and transcriptomics can help resolve this heterogeneity, we performed whole-genome sequencing (WGS) and RNA-seq on 20 tumors diagnosed as MPNST (**Supplementary Table S1, Supplementary Fig. S1**). We provided biological context at the transcriptional level, by including benign and premalignant lesions, derived Schwann cell primary cultures and established MPNST cell lines. Because transcription factors (TFs) capture lineage and cell-state identity, we focused on the expression of a curated set of 1,416 human TFs (34).

Unsupervised analysis of TF expression indicated that diagnosed MPNSTs do not form a single compact entity. Principal component analysis (PCA) and hierarchical clustering of the 20 primary tumors revealed two reproducible clusters, which we termed C1 and C2 (**Fig. 1A–C**). In the same TF expression space, benign neurofibromas and derived Schwann cell cultures were separated from malignant tumors, and primary tumors were separated from cultured cell lines **(Fig. 1D–E**), supporting that the TF-based embedding captures biologically meaningful axes of variation.

**Figure 1.**
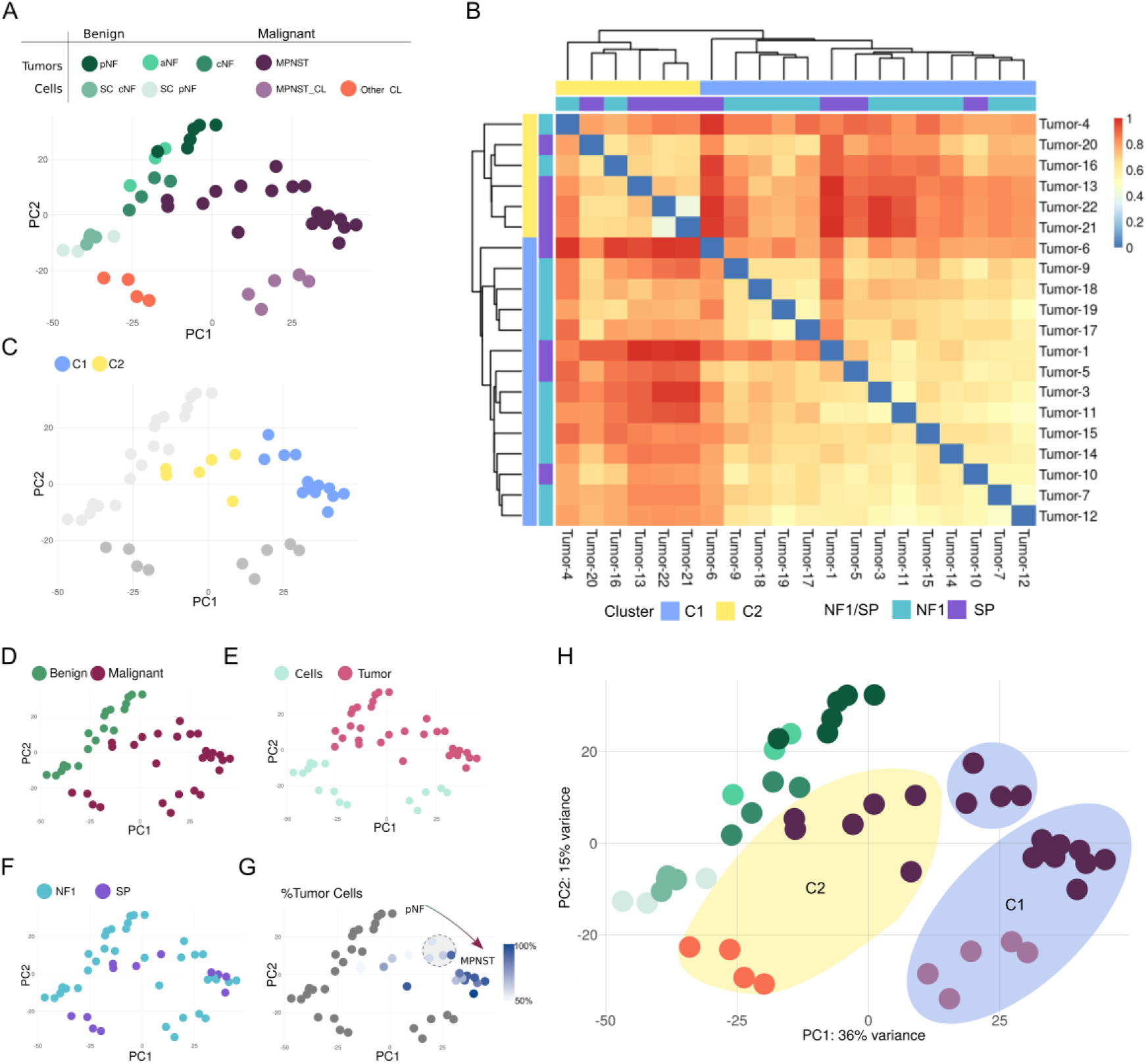
Transcription factor expression identifies two major clusters among tumors diagnosed as MPNST. **A)** Principal component analysis (PCA) of all samples analyzed by RNA-seq using the expression of 1,416 human transcription factors (TFs). **B)** Unsupervised hierarchical clustering of the 20 primary tumors diagnosed as MPNST using Euclidean distance on TF expression identifies two major groups, termed C1 and C2. **C)** PCA highlighting the distribution of the 20 primary tumors according to C1 and C2 cluster assignments. **D)** PCA showing the separation of benign and malignant samples. Benign samples include cNFs, pNFs, ANNUBPs and Schwann cell cultures derived from cNFs and pNFs. Malignant samples include tumors diagnosed as MPNST and previously characterized MPNST cell lines. **E)** PCA indicating the separation between primary tumors and cultured cells. **F)** PCA showing the distribution of NF1-associated and sporadic samples. **G)** PCA with primary malignant tumors colored according to the estimated tumor cell fraction derived from WGS **H)** PCA showing the position of previously characterized cell lines relative to the two transcriptional clusters. Genuine MPNST cell lines align with C1, whereas MPNST-mimicking cell lines align with C2.

A subset of NF1-associated tumors localized closer to benign samples along the pNF–MPNST axis and showed ∼50% tumor cell fraction estimated from WGS copy number analysis (**Fig. 1F-G**), consistent with substantial admixture of adjacent benign tissue. When we overlaid previously characterized MPNST cell lines (25), genuine MPNST lines aligned with C1, whereas MPNST-mimicking lines aligned with C2 (**Fig. 1H**). Together, these results defined two major transcriptional clusters among the 20 tumors diagnosed as MPNSTs and motivated a dedicated genomic dissection of the features distinguishing C1 from C2.

### MPNST clustering by TF expression is dominated by PRC2 status

Tumor suppressor gene (TSG) inactivation is a defining feature of MPNSTs. We performed an integrated analysis of copy-number variants (CNV), small nucleotide variants (SNV) and structural variants (SV) to obtain a comprehensive view of inactivation events affecting the most recurrently altered TSGs in MPNST (*NF1*, *CDKN2A*, and PRC2 components *SUZ12/EED*), as well as *TP53* **(Supplementary Table S2**).

Across TF-defined clusters, *NF1* and *CDKN2A* inactivation were more frequent in C1 than in C2 (**Fig. 2A-B**). *CDKN2A* was completely inactivated in all C1 tumors, and in half of these cases, the inactivation was mediated by translocations, as previously reported (25).

**Figure 2.**
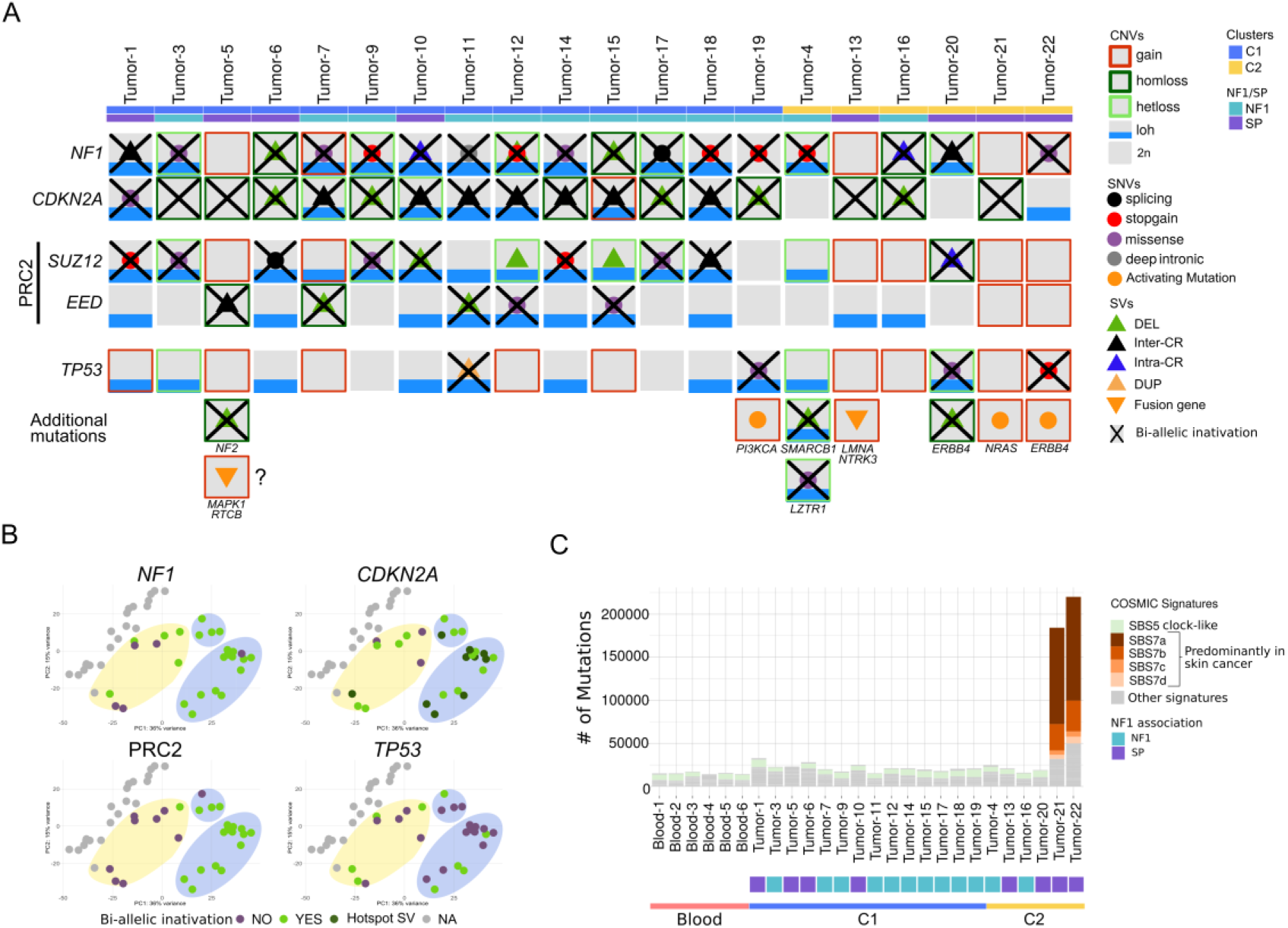
The transcriptional split between C1 and C2 is dominated by PRC2 inactivation. **A)** Integrated view of the genomic and mutational status of the main tumor suppressor genes recurrently altered in MPNST (*NF1*, *CDKN2A*, *SUZ12*, *EED*, and *TP53*) across the 20 tumors diagnosed as MPNST. Activating oncogenic mutations and fusion genes are also indicated. **B)** Principal component analysis (PCA) of transcription factor (TF) expression, colored according to the functional status of individual tumor suppressor genes. PRC2 inactivation is defined as the biallelic loss of *SUZ12* or *EED*. **C)** Total number of point mutations and COSMIC signatures composition of tumors diagnosed as MPNSTs and blood samples. Blue shadow represents C1, and yellow shadow is C2.

However, the most differential event between clusters was PRC2 inactivation: all but one C1 tumor showed biallelic inactivation of *SUZ12* or *EED*, whereas PRC2 inactivation was rare in C2 (1/6 tumors). Consistent with PRC2 being the main driver of the TF-based separation, we detected loss of H3K27me3 in genuine MPNST cell lines that co-clustered with C1, but not in the MPNST-mimicking cell lines that clustered with C2 **(Supplementary Fig. S2**). The frequency of *TP53* inactivation was low overall, particularly in C1. In fact, *TP53* alterations were significantly more frequent in MPNST cell lines (5/8) (17) than in primary tumors (4/20) (p=0.03), suggesting that *TP53* inactivation may facilitate adaptation to *in vitro* growth conditions (35).

Mutational features comprising small variants further distinguished the two clusters. C1 tumors displayed a mutational landscape comparable to blood controls, with low mutation burden and enrichment for the aging-associated clock signature SBS5 (**Fig. 2C**). The few C1 tumors harboring activating events lacked complete inactivation of at least one of the three major TSGs: Tumor-5 retained NF1 activity and carried a *MAPK1–RTCB* fusion gene, while Tumor-19, the only PRC2-wildtype tumor within C1, harbored an activating *PIK3CA* mutation (**Fig. 2A**). In contrast, C2 tumors retained PRC2 function and more frequently harbored activating events, including an *LMNA–NTRK1* fusion gene (Tumor-13) and mutations in *NRAS* (Tumor-21) and *ERBB4* (Tumor-22). Notably, Tumor-21 and Tumor-22 also exhibited markedly higher mutational burden and a strong enrichment for the UV-associated SBS7 signature, raising the possibility that a subset of C2 cases were in fact melanoma misclassified as MPNST (**Fig. 2A, 2C**).

### Translocation-mediated *CDKN2A* inactivation is already present in ANNUBP lesions

We previously identified translocations as a mechanism of *CDKN2A* inactivation in MPNSTs (25). Expanding this analysis to precursor lesions, we performed WGS on 7 ANNUBPs (**Supplementary Table S1**). Complete *CDKN2A* inactivation was present in all ANNUBPs, frequently mediated by translocations mapping to the same intronic hotspot observed in MPNSTs (**Fig. 3A–B; Supplementary Fig. S3**) (25). Furthermore, analysis of two distinct MPNSTs from the same patient revealed independent translocation breakpoints at this hotspot (**Fig. 3C**), underscoring the fragility of this locus in the cell of origin, at least in some patients. Notably, while SV burden was globally comparable between clusters, *CDKN2A* inactivation via translocations was exclusive to C1 (0/6 in C2) (**Fig. 3D–E**).

**Figure 3.**
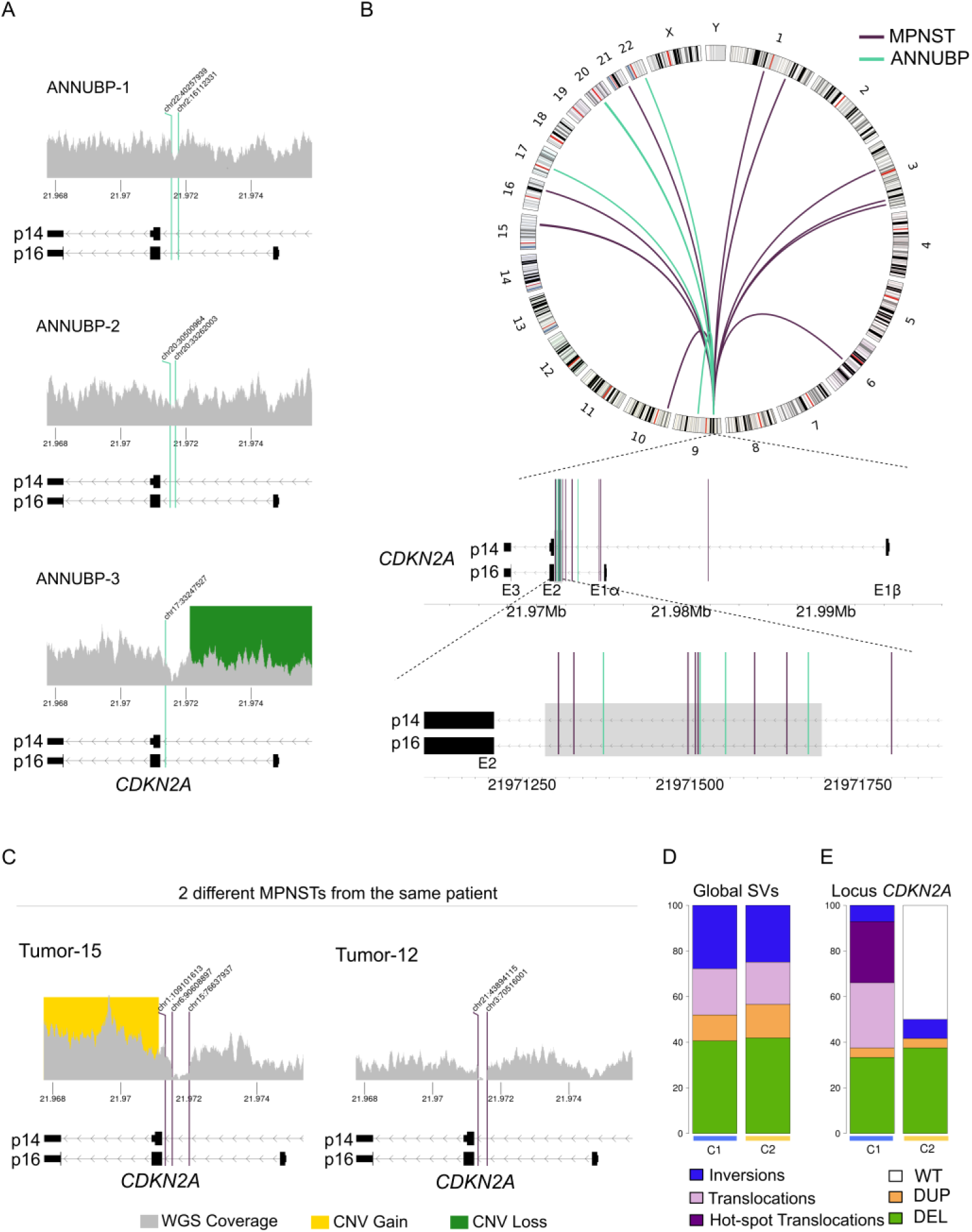
*CDKN2A* inactivation is recurrently mediated by translocations and is already present in ANNUBPs. **A)** Structural variants affecting the *CDKN2A* region in three ANNUBP samples analyzed by WGS. BAM coverage density, gene structure for p14 and p16 isoforms, rearrangements and deletions are displayed. **B)** Circos plot showing all translocations inactivating *CDKN2A* in MPNSTs and ANNUBPs with links to the partner regions. Below, exact position of all translocations inside the *CDKN2A* gene, with the lowest part highlighting those in the hotspot region. **C)** Structural variants affecting the *CDKN2A* region in two tumors from the same patient (Tumor-12 and Tumor-15). Gray shading indicates BAM coverage density, and vertical lines indicate breakpoint positions. **D)** Proportion of SVs of each type relative to the total number of SVs detected in tumors from the C1 and C2 clusters. DUP (duplication), DEL (deletion), WT (wildtype). **E)** Proportion of samples in each cluster (C1 and C2) harboring at least one SV affecting the *CDKN2A* locus ±1 mb

### MPNSTs from C1 share recurrent genomic features defining a three-stage model for MPNST tumorigenesis

While the genomes of all MPNSTs displayed hyperploidy and extensive rearrangement, a refined copy-number (CN) analysis —manually corrected for tumor purity and ploidy (see **Methods** and **Supplementary Fig. S4**)— uncovered genomic features exclusive to C1 tumors: near tetraploidy and recurrent large regions of copy-neutral loss-of-heterozygosity (CN-LOH) (**Fig. 4A**; **Supplementary Fig. S5**). C1 genomes showed a significant enrichment for even copy-number states (**Supplementary Fig. S5A**), and we identified recurrent CN-LOH affecting specific chromosomes or chromosome arms (1p, 4, 9p, 10, 11, 14q, 16q, 17) in over 50% of C1 cases (**Fig. 4A and Supplementary Fig. S5**). Notably, without the applied ploidy and purity corrections, most of these regions would have been misclassified as genomic losses (**Supplementary Fig. S4**). To confirm that these regions were *bona fide* 2n copies rather than deletions, we validated the CN-LOH calls on chromosomes 1 and 11 in three independent MPNST cell lines (sNF96.2, NF90-8, and ST88-14) (**Fig. 4B**). Using fluorescence in situ hybridization (FISH) with probes located on both arms of each chromosome (**Fig. 4C, Supplementary Table S3**), we demonstrated that these CN-LOH regions were indeed diploid (2n) within a tetraploid context (**Fig. 4B–C**). This pattern is consistent with a diploid precursor clone acquiring chromosome-specific losses followed by a whole-genome doubling (WGD) event (**Fig. 4D**) that contributes to the malignant progression.

**Figure 4.**
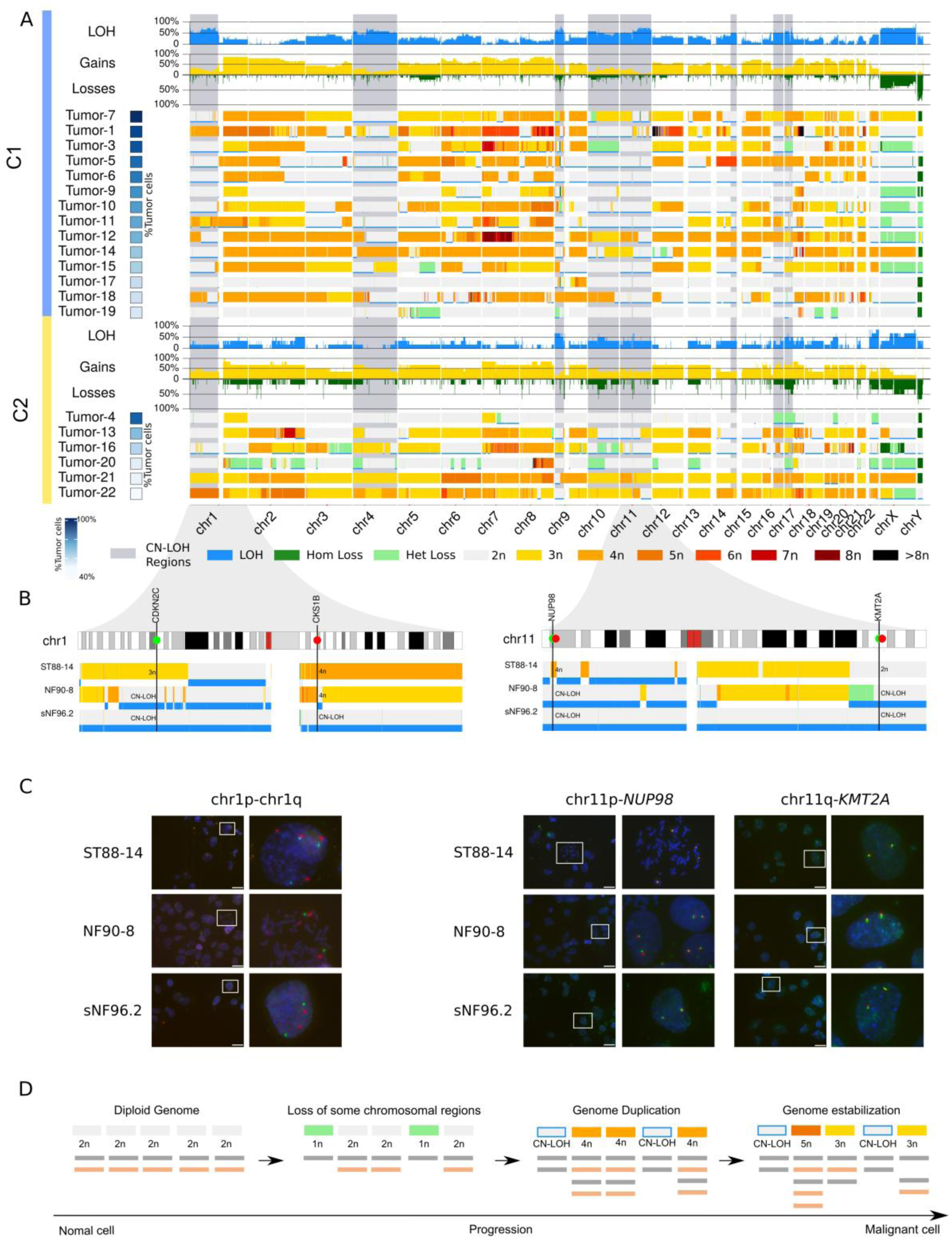
C1 tumors show recurrent copy-neutral loss of heterozygosity regions and a characteristic copy-number profile. **A)** Copy-number and loss-of-heterozygosity (LOH) profiles of tumors from the C1 and C2 clusters after adjustment for tumor purity and ploidy. For each cluster, the lower section shows the copy-number and LOH profile of individual tumors, ordered by estimated tumor cell fraction. Gains are shown in warm colors and losses in green, with darker shades indicating higher-amplitude alterations. LOH is indicated by a blue track below the copy-number profile. The upper section summarizes the accumulation of gains, losses, and LOH events across tumors in each cluster. Gray vertical bands indicate recurrent copy-neutral LOH (CN-LOH) regions observed in more than 50% of C1 tumors. **B)** Schematic representation of chromosomes 1 and 11 showing the positions of the probes used for FISH validation in ST88-14, NF90-8, and sNF96.2 cells, together with the copy-number state of each locus in the corresponding samples. **C)** Representative FISH images of the loci shown in B in the indicated cell lines. **D)** Schematic model summarizing a sequence of genomic events that could lead to the genomic profiles we see in C1.

Our analysis so far revealed, in C1, a consistent set of inactivated TSGs, with *NF1* and *CDKN2A* already inactive in ANNUBPs, and PRC2 only in MPNSTs and a set of recurrent genomic features compatible with a common progression path. This lends itself to a simple three-stage model for MPNST tumorigenesis: first, the inactivation of TSGs, partially happening in premalignant lesions (Initiation), followed by an extensive genomic rearrangement (Progression) and a final genome stabilization and adaptation to the altered malignant state (Stabilization) (**Supplementary Fig. S5E**).

### Dedicated genomic re-analysis of an external dataset confirms the identified MPNST genomic hallmarks

To assess the generalizability of our findings, we examined an independent cohort of 75 MPNSTs previously characterized by the GeM consortium (18). To ensure comparability with our samples and reduce batch effects, we performed a complete re-analysis of the raw WGS and RNA-seq data, applying the same dedicated analysis pipeline used for our cohort. Before the analysis, we first filtered the GeM samples based on data quality and tumor purity, resulting in a set of 50 high-quality MPNSTs (31 NF1-associated and 19 sporadic) and 27 normal samples (**Supplementary Fig. S6-S7, Supplementary Table S4**). The re-analysis proved crucial, as we identified additional genetic and genomic alterations in 25% of the samples, like the identification of unreported inactivating mutations in *NF1*, *CDKN2A*, *EED,* or *TP53*, the presence of fusion genes or oncogenic variants in several samples (**Supplementary Table S5 and S6**).

Projecting the GeM transcriptomes onto our original transcription factor PCA space revealed a distribution highly consistent with our initial set of MPNSTs **(Fig. 5A**): 27 tumors clustered with C1 cluster, 19 clustered with C2, and 4 located in between (**Fig. 5A-B**).

**Figure 5.**
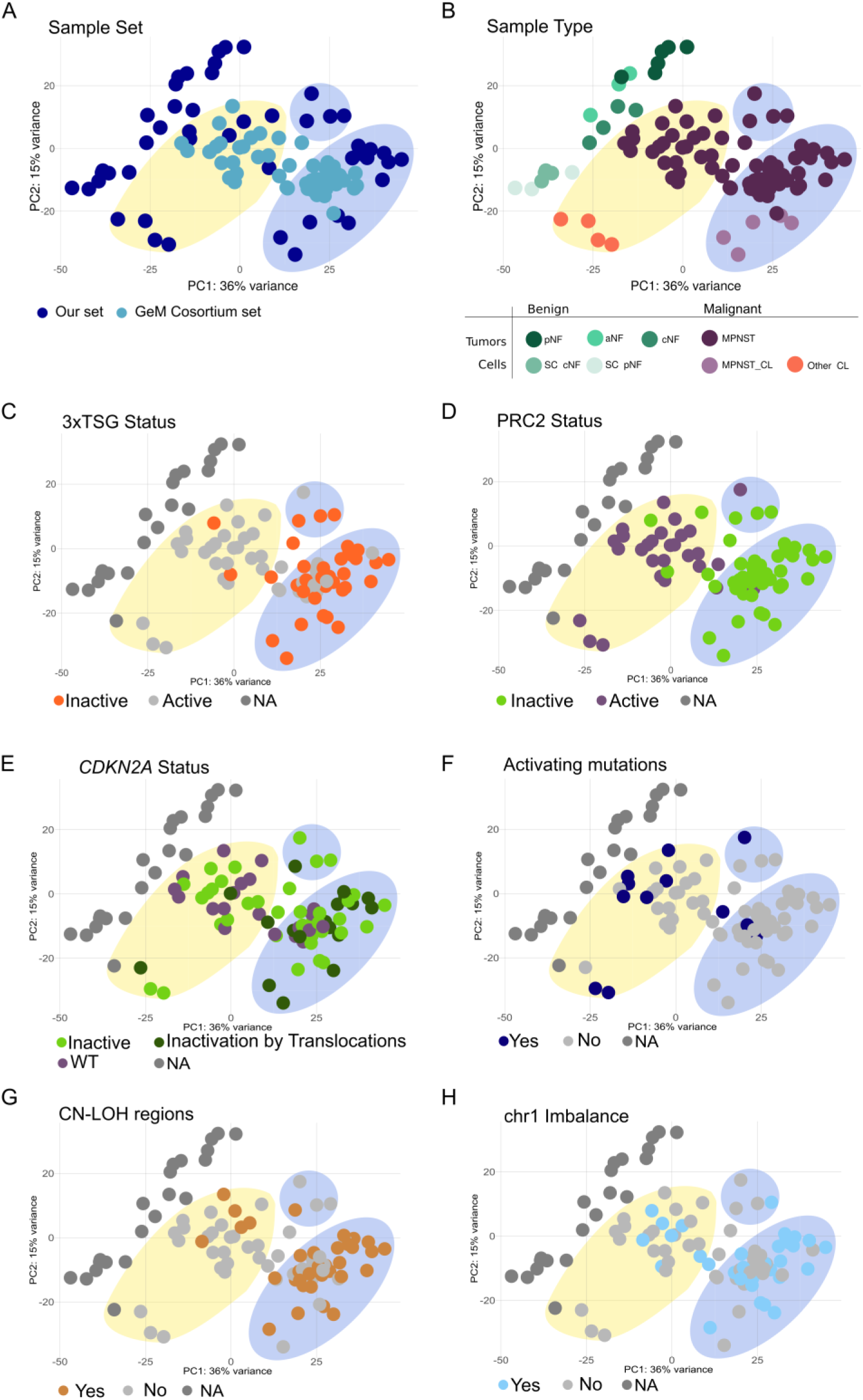
Genomic hallmarks identified in the in-house cohort are validated in the GeM Consortium dataset. **A)** Principal component analysis (PCA) of transcription factor (TF) expression showing the *in-house* cohort and the projection of the GeM Consortium samples according to sample provenance. Blue shading indicates the C1 region and yellow shading indicates the C2 region. **B)** PCA showing the distribution of samples according to sample type. **C)** PCA showing the distribution of samples according to the combined inactivation status of *NF1*, *CDKN2A*, and PRC2. **D)** PCA showing the distribution of samples according to PRC2 status. **E)** PCA showing the distribution of samples according to *CDKN2A* status. **F)** PCA showing the distribution of samples according to the presence or absence of activating oncogenic alterations. **G)** PCA showing the distribution of samples according to the presence of CN-LOH in more than 50% of the previously identified recurrent C1-associated CN-LOH regions. **H)** PCA showing the distribution of samples according to the presence or absence of chromosome 1 imbalance.

The combined analysis of both cohorts validated the genomic hallmarks identified in our samples. The distribution of TSG inactivation was conserved and clustering was again dominated by PRC2 status (**Fig. 5C–D**). We confirmed that, within C1 tumors, *CDKN2A* inactivation was predominantly mediated by translocations (**Fig. 5E**). We also confirmed that C1 tumors consistently lacked activating mutations (**Fig. 5F**) and recapitulated the specific CN-LOH signature (**Fig. 5G**) and the chromosome 1 copy-number imbalance (**Fig. 5H**) observed in the initial set of samples.

### From two clusters to three genomic groups based on TSG inactivation

We wanted to explore going beyond the TF-based clusters C1 and C2, which were dominated by the profound effect of PRC2 inactivation on transcriptional patterns (36). To do so, we turned to the initial step of the proposed three-stage MPNST model of tumorigenesis: the tumor suppressor gene inactivation.

When we classified MPNSTs according to the combination of inactivated TSGs present in each tumor, three clear major groups emerged: the first one, G1 (n=32), with *NF1*, *CDKN2A* and PRC2 inactivated; the second, G2 (n=10), with *NF1, TP53* and PRC2 inactivated but *CDKN2A* active; and the third, G3 (n=8), with *NF1* and *CDKN2A* inactive and PRC2 active (**Fig. 6A, Supplementary Table S7**). We grouped all other less frequent TSG combinations and all samples with activating mutations in a fourth group called “Others” (n=20). Although these groups were defined mainly by the status of TSG inactivation, they correlated surprisingly well with other genetic, genomic, histological and clinical features (**Fig. 6A, Supplementary Table S7**). G1 MPNSTs were characterized by translocation-mediated inactivation of *CDKN2A* at the identified hotspot region, bearing almost no additional mutations in other TSGs like *TP53, RB1* and *PTEN*. They had a hyperploid genome, exhibiting characteristic CN-LOH regions and copy number imbalance in chromosome 1 (**Supplementary Fig. S8**), and were associated with a conventional MPNST histology. MPNST in G2 bore non-functional PRC2, mainly caused by mutations in *EED*, in contrast to G1 tumors, which were mainly caused by mutations in *SUZ12.* All but one MPNST had a complete inactivation of *TP53,* a proportion significantly larger than in any other group. Importantly, most of them displayed a diploid genome with generalized CN-LOH (**Supplementary Fig. S8 and S9**) and strikingly, all but one developed in males. On the other hand, G2 MPNSTs were associated with a histology described as conventional MPNST with heterologous elements, frequently with Rhabdomyoblastic traits (**Supplementary Table S5 and S7**). As for G3 MPNSTs, their genomes were hyperploid and had more structural variant breakpoints than any other group. However, in contrast to G1, they had no CN-LOH in the identified specific regions nor chromosome 1 imbalance. It was associated with conventional MPNST histology and contained only NF1-associated cases. Finally, the group “Others” concentrated sporadic cases and was not enriched in any specific histology class. It was diverse in most other metrics as well, and contained at least some tumors with specific features that suggested they were not true MPNSTs but misclassified tumors (**Supplementary File S1).**

**Figure 6.**
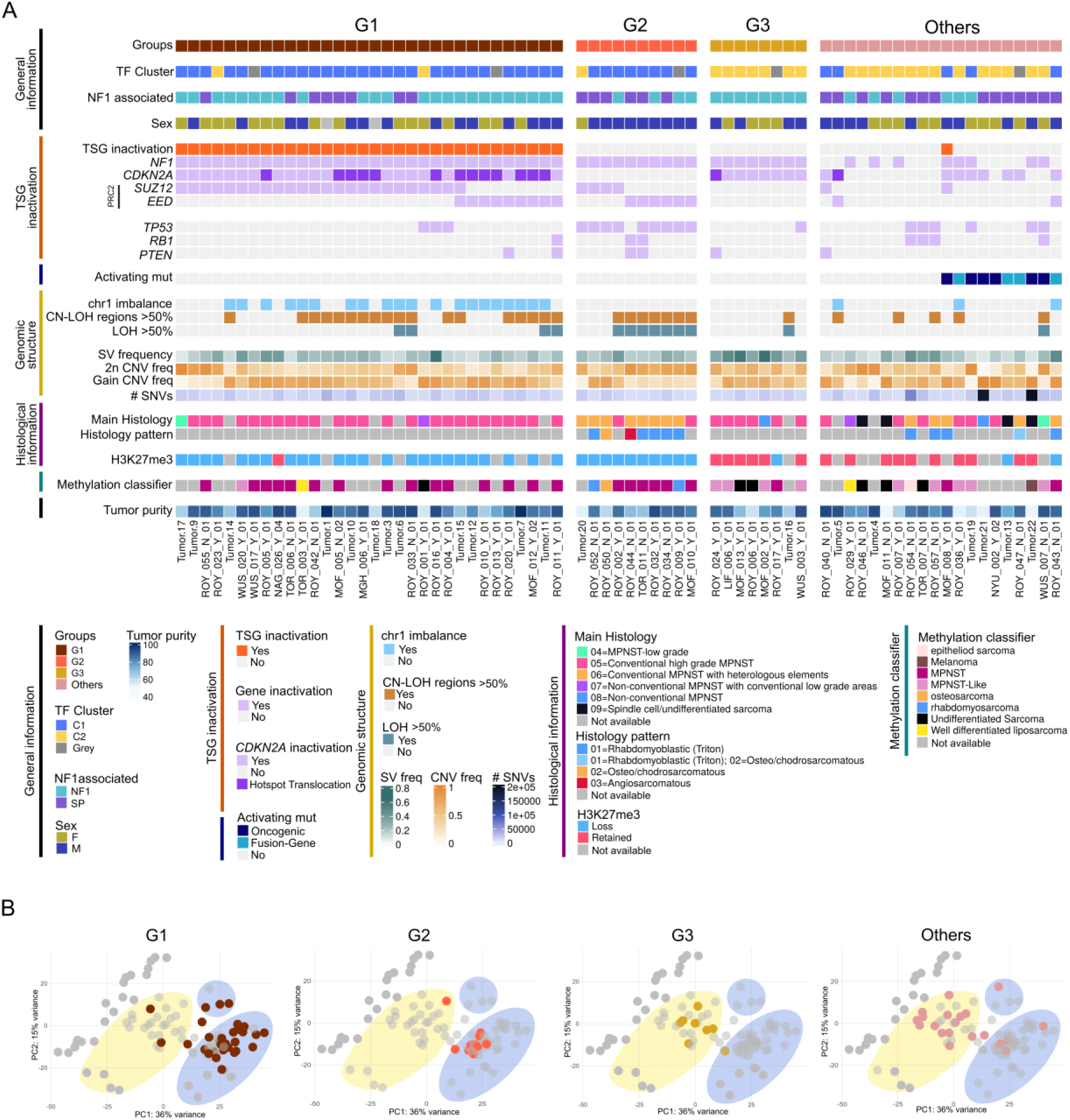
Three major genomic groups emerge from classification by tumor suppressor gene inactivation status. **A)** Summary of the 70 tumors analyzed across the combined cohort, grouped into G1, G2, G3, and “Others” according to the status of the main recurrently altered tumor suppressor genes (*NF1*, *CDKN2A*, PRC2, and *TP53*) and the presence of activating oncogenic alterations. Clinical, genetic, genomic and histological information from each tumor is also included. **B)** PCAs depicting the distribution of all MPNST samples from each group.

Although gene expression profiles played no part in defining these groups, the strong impact of PRC2 inactivation on transcriptome partially linked the TSG-defined groups to the TF-based clusters (**Fig. 6B**), with G1 and G2 concentrating most samples in C1 and G3, and “Others” concentrating most samples in C2. The impact of PRC2 inactivation was also visible in the methylome classifier, with samples in G1 and G2 classified mostly as MPNSTs and G3 as MPNST-like (**Fig. 6A**). The methylation-based classification of samples in the “Others” group was again diverse.

### Each MPNST progression path is linked to the initial TSG inactivation

Each genomic group was not only defined by the TSG inactivating signature, but enriched for specific patterns of ploidy, LOH, copy-number and structural alterations. Placed in the framework of our three-stage model of MPNST development, these genomic features defined distinct progression trajectories for each MPNST group. The trajectories comprised different TSG inactivation patterns, followed by specific mechanisms of genome rearrangement, and a particular stabilization into characteristic copy-number profiles **(Fig. 7A–B; Supplementary Fig. S10**). G1 tumors present a hyperploid genome, recurrent CN-LOH at specific regions, chromosome 1 imbalance and gains in chromosomes 2, 7 and 8 (**Fig. 7A-B**). This genomic state is compatible with a trajectory initiated by the inactivation of *NF1, CDKN2A* and PRC2, followed by hemizygous loss of parts of the genome, including 1p, a whole-genome duplication event and a few additional genomic alterations resulting in a stable nearly tetraploid genome. G2 tumors, in contrast, present largely diploid genomes, with extensive genome-wide LOH and a recurrently gained chromosome 8 (**Fig. 7A-B; Supplementary Fig. S8 and S9**). This is compatible with an initial loss of *NF1, TP53* and *EED,* then the loss of a near-complete set of chromosomes, excluding chromosome 8, followed by endoreduplication of the remaining set and a stabilization into a diploid LOH–dominated genomic state. G3 tumors are characterized by high structural-variant burden (**Fig. 6A and Fig. 7Aii**), extensive rearrangements, and recurrent gains on chromosomes 2p, 7, and 12 (**Fig. 7A–B**). These characteristics are consistent with an initial loss of only *NF1* and *CDKN2A*, followed by a highly catastrophic genomic event, converging on a stabilized state that differs from G1 and G2, especially in the lack of a high frequency of gained chromosome 8 (**Fig.7A-B**). Copy-number differences among G1–G3 could potentially aid the diagnosis and classification of MPNSTs (**Fig. 7B**).

**Figure 7.**
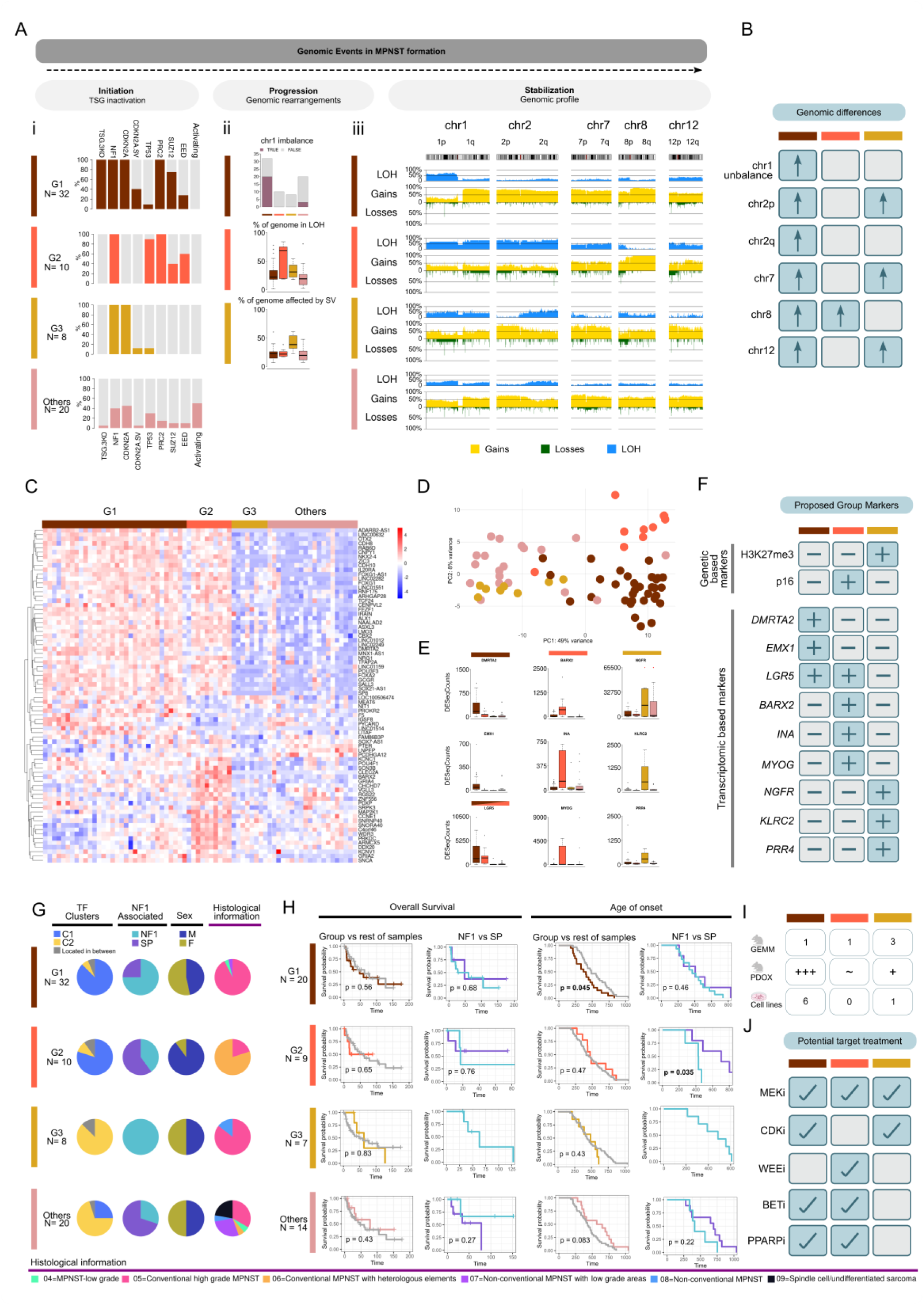
TSG-defined groups are linked to different progression paths and show clinical differences. **A)** Genomic features of the different groups. Ai, Percentage of samples in each group bearing the inactivation of key TSGs and activating mutations. Aii, Frequencies of chr1 imbalance, percentage of genome in LOH, and percentage of genome affected by an SV across groups. Aiii, Frequency of gains, losses and LOH for each group over a selected chromosome (chr1, chr2, chr7, chr8, chr12). **B)** Summary of the copy-number status of key genomic regions across groups. **C)** Heatmap showing the expression of the 76-gene signature across the 70 tumors. **D)** PCA representing the distribution of the 70 tumors according to the expression of the 76-gene signature. **E)** Gene expression boxplot of potential markers differentiating between groups. Red dots represent outliers. **F)** Summary of a selection of potential gene markers to differentiate between MPNST groups. **G)** Pie charts representing the relation between groups and molecular, clinical and histopathological features: TF-based clusters (C1, C2, or Grey), sporadic (SP) or NF1-related (NF1), sex (male (M), female (F)) and main histological classification. **H)** Kaplan-Meier curves representing overall survival and age of onset across groups. For each a plot comparing the group versus the rest of the samples and NF1 versus SP cases inside the group. **I)** Quantity of GEMM, PDOX and cellular models available for each group. **J)** Potential target and personalized treatment for each group.

### Each MPNST group expresses specific gene signatures

To dissect the transcriptional landscape underlying genomic subgroups, we performed a differential expression analysis across G1-G3, identifying broad transcriptional differences between groups. We applied a feature selection approach based on Boruta followed by recursive feature elimination with cross-validation (RFECV), which produced a 76-gene signature that separated genomic subtypes with good performance in cross-validation (≈83% accuracy) (**Fig. 7C–D**,). Gene Set Enrichment Analysis (GSEA) comparing PRC2-inactive (G1 and G2) versus PRC2-active (G3) subgroups revealed a strong link to early developmental programs in the PRC2-deficient tumors, consistent with the loss of epigenetic repression (**Supplementary Fig. S11A**) (36). Beyond this shared axis, group-specific programs were also present. In particular, differences between the PRC2-deficient subgroups (G1 and G2) showed a significant enrichment in muscle-related terms in G2 (**Supplementary Fig. S11B**), providing a transcriptomic basis for the rhabdomyoblastic (“Triton”) differentiation frequently observed at the histological level in G2. In addition to a gene expression signature that could act as an MPNST group classifier, we aimed to identify specific immunohistochemical (IHC) analysis-based markers to potentially aid the diagnosis of MPNST groups. Based on the differential expression at group level, at individual sample level and regarding the availability of IHC assays, we manually curated a list of 1703 genes and selected 9 markers with strong group specificity: 2 expressed in G1 (*EMX1*, *DMRTA2*); 1 expressed in G1 and G2 (LGR5); 3 expressed in G2 (*INA*, *BARX2*, *MYOG*) and 3 expressed in G3 (*KLRC2*, *NGFR*, *PRR4*) (**Fig. 7E**). These markers, together with existing ones (H3K27me3, p16), might be used in a differential diagnosis of MPNST groups (**Fig. 7F**) (**Supplementary Fig. S10**).

### Preclinical and clinical impact of MPNST genomic groups

Genomic subgroups were also associated with distinct clinical features (**Fig. 7G**). G1 and G3 contained mostly NF1-associated cases (100% NF1-associated in G3) and G2 and “Others” were enriched in sporadic cases. Sex was evenly split for all groups except for G2, which showed a strong male predominance (9M, 1F). Although cohort size limited the statistical power of the survival analysis, we detected group-wise differences in the age of onset (**Fig. 7H),** including within G2 when stratified by etiology (NF1-associated versus sporadic). Genomic groups also had uneven availability of preclinical animal models (**Fig. 7I**). Three genetically engineered mouse models (GEMM) exist, representing G3 (37–39), while only one GEMM recapitulates G2 (26) and G1 (40). In the case of MPNST PDOX models, there exist models for the three groups, although G1 is by far the most represented (35,41,42). Regarding cellular models, out of 7 MPNST cell lines for which we had complete WGS data, 6 presented the G1 (17,35,43) and 1 represented G3 (35). Finally, given the different set of initial TSG inactivation, personalized medicine strategies (36,42) may need to be tailored to the different groups (**Fig. 7J**) and move beyond treating MPNSTs as a single entity.

## DISCUSSION

MPNST is a biologically heterogeneous and genomically complex sarcoma, and this heterogeneity, together with overlapping histological presentations with other tumor entities, has long complicated an accurate diagnosis and hindered the development of effective personalized treatment strategies (11,13,17,44). A consensus representation of an MPNST genome was missing, as well as a basic unifying model of MPNST formation. Knowing the genomic complexity of MPNSTs, we used a careful bioinformatic analysis of WGS data to cover the spectrum of potential genomic alterations and mutational mechanisms involved in MPNST formation (17,25) and completely characterize, first, the genome of our own set of 20 MPNSTs and 7 ANNUBPs, and later 50 MPNSTs from the GeM Consortium (18) selected after a stringent quality control. Performing a careful and dedicated bioinformatic analysis was key to getting a complete genomic picture from both sample sets. This complete data naturally unfolded into a simple three phase model of MPNST development: 1) an initiation phase, characterized by the loss of specific combinations of TSGs; 2) followed by a progression phase in which the genome is catastrophically reorganized; and 3) finally, a stabilization phase, resulting in a viable tumor cell, with a specific identity and a genomic copy-number alteration (CNA) profile. Our transcriptomic analysis quickly showed that global gene expression in MPNSTs is strongly shaped by PRC2 status, mirroring the pattern observed in the methylome (30). “Thus, when classified according to expression or methylation profiles, MPNSTs separate into two major clusters, making additional subgroup structure difficult to resolve. Genomic characterization provided an additional layer of information that allowed, for instance, to separate MPNSTs from Malignant triton tumors (MTTs), something not possible by methylome classification (30).

Another relevant genomic finding is that CDKN2A inactivation caused by translocations, previously identified in MPNSTs (18,25), is already present in ANNUBPs, highlighting this inactivation mechanism as preponderant in the cells originating these pre-malignant lesions. In some patients, distinct lesions harbored independent translocations, suggesting that fragility at the CDKN2A locus may represent a risk factor for MPNST development.

Furthermore, we also discovered the existence of highly recurrent CN-LOH regions in G1 MPNSTs, the largest MPNST group. These regions were previously characterized as recurrent “genomic losses” (13,18,33,45,46). This was due to the inherent limitations, technological and biological, of the techniques and bioinformatic programs used for assessing CNAs in hyperploid genomes such as those of MPNSTs, underestimating copy-number gains and overestimating copy-number losses (47,48). Accounting for the presence of non-tumoral cells before performing different CNA estimates, considering distinct ploidy scenarios and combining genomic tools with FISH analysis were instrumental for their identification and validation. The existence of these CN-LOH regions together with a large proportion of the genome in even copy-number, supports a model in which these regions are lost in heterozygosity before a genome duplication event takes place (49). Finally, our analysis corroborated the low mutation burden of MPNSTs compared to other tumors (16,17,50) as well as their generalized lack of gain-of-function mutations, with the exception of NTRK-related MPNSTs (see below).

MPNST heterogeneity is well documented, particularly concerning histological presentations (11,51). There also exist other tumor entities that mimic these histological presentations, and there is a clear lack of biomarkers for a precise differential diagnosis (11). Echoing this heterogeneity, different works performing thorough molecular analyses identified groups of MPNSTs with different profiles (18,30,33,52,53). However, we still needed to integrate all these molecular findings and gain a comprehensive general view of MPNST heterogeneity. In the present work, classifying the 70 MPNSTs just by their combination of inactivated TSGs, that is, by its first developmental step, astonishingly uncovered three homogeneous MPNST groups and one additional group of other miscellaneous entities. Most importantly, each identified MPNST group was characterized by a distinct progression path and by a distinct genomic structure, histology, expression profile and potential clinical behavior (**Supplementary Fig. S10).** TSG inactivation constitutes the first step of MPNST development. The pattern of TSG inactivation is not completely understood, but it might reflect a different physiology and/or identity of the cell originating each MPNST group, or, conversely, be just a matter of chance. G1 tumors lose *NF1*, *CDKN2A,* and PRC2, in this order, as evidenced by the progression pNF-ANNUBP-MPNST. We speculate that G2 MPNSTs lose first *NF1*, then *TP53* and finally PRC2, since, at least in iPSCs, the loss of PRC2 in an *NF1*(-/-) iPSCs is not viable if *CDKN2A* is WT (36). We presume that prior loss of *TP53* may relax this constraint. In contrast, G3 tumors show loss of *NF1* and *CDKN2A* only, together with extensive SV-driven genome rearrangement, similar to spontaneous progression in *Prss56-CRE* mice (54).

Finally, the group termed “Others” contains different miscellaneous entities (**Supplementary File 1)**. For instance, it contains rare MPNSTs bearing *NTRK* fusions (55–57), each exhibiting a high expression of the respective *NTRK* gene involved, supporting the use of a pan-TRK IHC assay. It also contains tumors with *FUS-TFCP2* fusion genes described in spindle cell rhabdomyosarcoma (58). In these cases, these fusion genes might also be key for their first initiation step instead of the loss of TSGs. The “Others” group may also contain other entities like melanomas, due to the presence of a high number of SNVs and associated mutational signatures in some tumors. However, we cannot discard the possibility that this group also contains misclassified genuine MPNSTs with distinct TSG inactivation signatures.

These TSG losses predispose tumor cells to undergo a genome reorganization through a disruptive genomic event: normally, a genome duplication in G1; in G2, commonly a loss of one chromosome complement followed by a reduplication resulting in generalized 2n in LOH (also in some G1 tumors); a genome duplication followed by a extensive genomic rearrangement in G3. This genomic rewiring needs to be resolved in a viable MPNST cell that bears a specific CNA profile, distinctive in each MPNST group. All these genomic differences, TSG inactivation profile, genome structure and CNA profile, presence of oncogenic mutations of fusion genes, could be used as biomarkers to complement pathological diagnosis.

G1 and G3 tumors associate with a conventional or classic MPNST histology, and G2 MPNSTs with entities containing rhabdomyoblastic differentiation, like MTT (30). Despite these histological associations, we also identified distinct expression profiles for each MPNST group, useful for building an MPNST group classifier. Several genes with a strong group specificity can be detected by immunohistochemistry and could serve as differential diagnostic biomarkers among MPNST groups. Some differentially expressed genes in G2 are muscle-related genes, consistent with its histological presentation.

G1 MPNSTs constitute around 65% of all MPNSTs, with a higher proportion of NF1 vs sporadic presentations. G2 accounts for 20% of MPNSTs and occurs almost exclusively in males. G3 accounts for 16% of MPNSTs and is composed exclusively of NF1-associated tumors. Overall survival trends of different MPNST groups were limited by sample size, although the age of onset was indeed different. Finally, there are differences regarding the existing *in vitro* and *in vivo* models representing the different MPNST groups. For instance, while different GEMMs exist for the less frequent G2 and G3 subtypes (26,37,38,54,59), there is a notable lack of *in vivo* models for the most abundant G1 (60). However, the opposite scenario occurs for MPNST PDOX mouse models.

A much wider, complete analysis of MPNSTs, ideally involving multiple sites in an international effort, could bring a definitive picture of the biological and clinical implications that these identified MPNST groups might hold.

## MATERIALS AND METHODS

### Human Sample Collection

Twenty primary tumors diagnosed as MPNSTs were collected from different Spanish hospitals: Hospital Sant Joan de Deu (HSJD) (n = 8), Hospital Universitari Bellvitge (HUB) (n = 6), Hospital Universitari Vall d’Hebron (HUVH) (n = 3), Hospital Universitari Germans Trias i Pujol (HUGTiP) (n = 2), Hospital Universitario la Paz (HULP) (n = 1). Besides, seven ANNUBPs were also collected from different Spanish hospitals: HSJD (n = 4), and HUB (n = 3). Clinical and pathology reports of each participant were collected when available (**Supplementary Table S8**). PNFs from the IGTP Hereditary Cancer sample collection (ISCIII: C.0002242), and derived primary cell cultures were also included. All patients provided written informed consent. The project was approved by the Clinical Research Ethics Committee of Hospital Universitari Germans Trias i Pujol (Badalona, Spain).

### Cell lines

In this study, we used six NF1-associated MPNST cell lines: S462 (RRID:CVCL_1Y70) (61), ST88-14 (RRID:CVCL_8916) (62), NF90-8 (RRID:CVCL_1B47) (63), sNF96.2 (RRID:CVCL_K281) (64), NMS-2 (RRID:CVCL_4662) (65), and three sporadic MPNST cell lines: STS-26T (RRID:CVCL_8917) (66), HS-Sch-2 (RRID:CVCL_8718) (67) and HS-PSS (RRID:CVCL_8717). Cells were grown in DMEM supplemented with 10% FBS (Gibco), 1x GlutaMAX (Gibco) and 500 U/mL Penicillin/500 mg/mL Streptomycin (Gibco) and maintained at 37°C under a 5% CO2 atmosphere. All cell lines were authenticated (17) and tested for mycoplasma.

### Nucleic acids extraction

Genomic DNA from tumors was extracted using the Gentra Puregene Kit (Qiagen) following the manufacturer’s instructions, after homogenization with a TissueLyser (Qiagen). Genomic DNA from cells was extracted using Promega Maxwell 16 instrument (Promega) according to the manufacturer’s instructions. All DNA was quantified using a Qubit fluorometer (Life Technologies) and its quality was assessed with Agilent 2200 TapeStation (Agilent).

Total RNA was extracted using TriPure Isolation Reagent (Roche) following the manufacturer’s instructions from flash-frozen tumors thawed in DMEM supplemented with 10% FBS and homogenized using a TissueRuptor II (Qiagen). RNA was quantified with a NanoDrop 1000 spectrophotometer (Thermo Fisher Scientific) and quality was assessed with Agilent 2200 TapeStation (Agilent).

### Immunocytochemical analysis

Cells were fixed in 4% paraformaldehyde (PFA) (Santa Cruz Animal Health) in PBS for 15 min at room temperature, permeabilized with 0.1% Triton X-100 in PBS for 10 min, blocked in 10% FBS in PBS for 15 min, and incubated with the primary antibody, H3K27me3 Rabbit mAb (RRID:AB_2616029) overnight at 4°C. An Alexa Fluor 568 goat anti-rabbit (Thermo Fisher Scientific) secondary antibody was used. Nuclei were stained with DAPI (Stem Cell Technologies, 1:1000). Slides were mounted with Vectashield (Vector Laboratories), and coverslips were secured with nail polish. Images were acquired using a DMI 6000B microscope (Leica) and LAS X software (Leica).

### CN-LOH regions validation by FISH

FISH was performed on ST88-14, NF90-8 and sNF96.2 cell lines following Metasystems protocols. The nuclei were fixed in Carnoy’s solution. The sample and probe were co-denatured by heating slides on a hotplate at 75°C (±1°C) for 2 min. Hybridization was carried out with 5 µL of probe, followed by overnight incubation in a humidified chamber at 37°C (±1°C). Post-hybridization washes consisted of immersion in 0.4× SSC (pH 7.0) at 72°C (±1°C) for 2 min, followed by washing in 2× SSC containing 0.05% Tween-20 (pH 7.0) at room temperature for 30 s. Slides were briefly rinsed in distilled water to prevent crystal formation, air-dried, counterstained with DAPI antifade, and examined under a fluorescence microscope. FISH analyses were performed on the Metafer FISH imaging system (MetaSystems). The following probes were used for FISH: XL CDKN2C/CKS1B, XL NUP98, XL KMT2A BA (MetaSystems). FISH capturing was performed in a Metafer Slide Scanning System (MetaSystems) with a Zeiss Axio Imager epifluorescence microscope equipped with a motorized stage, 10× and 63× oil plan APOCHROMAT objectives and specific filters for DAPI, Spectrum Green and Spectrum Orange (Nikon).

### Whole Transcriptome Sequencing (RNA-seq)

RNA-seq data of 41 samples were previously generated in different studies (17,36,68,69). RNA-seq libraries of five additional samples sequenced for this work were prepared at BGI (Shenzhen, China) using DNBSEQ standard protocols (**Supplementary Table 1)**.

### RNA-seq analysis

Transcript-level abundance from RNA-seq data was estimated with Salmon v1.8.0 (70) using UCSC RefSeq (refMrna) transcript annotations and the hg38 reference genome. These estimates were then imported into R and summarized to gene-level expression matrices using the tximport R package (71). After that, we kept only genes with more than five counts in three or more samples.

For TF-based analyses, we restricted the expression matrix to a curated set of 1,416 human transcription factors (TFs) (34). Unsupervised clustering was performed on the 20 primary MPNST tumors from our in-house cohort using Euclidean distance. PCA was performed using all samples in the in-house dataset. GeM whole-transcriptome data were processed using the same pipeline and projected onto the in-house PCA using the PCA loadings.

### Whole-genome sequencing (WGS)

WGS data for nine tumors were generated in a previous study (25) (**Supplementary Table 1)**. Whole-genome sequencing for the remaining eleven tumors and five normal samples was performed for this study at BGI (Shenzhen, China). Libraries were prepared following standard DNBseq protocols and sequenced on a BGISEQ-500 (paired-end, 2×150 bp).

### Additional samples from the GeM Consortium

We accessed WGS and RNA-seq data from the GeM Consortium (18) (EGAD00001008608) to extend our local cohort. We filtered cases to select complete, high-quality samples. Specifically, we required (i) availability of WGS and RNA-seq data and reported results from the methylation classifier, (ii) consortium-reported tumor purity >50%, and (iii) for patients with multiple tumor samples, selection of the purest sample. We then re-estimated tumor purity using the same procedure applied to our in-house cohort and excluded samples with estimated purity <50%. Quality control of the associated normal samples revealed problems and inconsistencies in several cases (**Supplementary Figs. 6 and 7; Supplementary Table 3**), resulting in a final GeM subset of 50 tumors and 27 normal samples.

### Variant Calling from WGS

Raw sequencing data were mapped with BWA-MEM (72) to the GRCh38.p14 reference genome. Because matched normal samples were not available for all tumors, SNVs were called with Strelka2 (73) in germline mode. Identified variants were then annotated with ANNOVAR (74) including population frequencies, clinical annotations and pathogenicity predictions.

To identify potentially pathogenic variants, we filtered annotated variants as follows: we selected exonic and splicing variants and removed all synonymous variants. Then, we filtered out variants with a population frequency (AF_popmax) > 1%, classified as benign in ClinVar (75), annotated as benign or likely benign in InterVar (76) or observed in more than one individual in our cohort. We then kept variants predicted damaging by ≥5/7 *in-silico* predictors (SIFT (77), PolyPhen2 HDIV (78), LRT (79), Mutation Taster (80), Mutation Assessor (81), FATHMM (82) and CLNSIG (75). Then, we filtered out those variants with a variant allele frequency (VAF) lower than 0.1, as these variants were deemed as unlikely to be present in the original malignant cell population. Finally, we removed variants in highly variable genes (*MUC3A*, *MUC5AC*, *OR52E5*, *OR52L1*, *SMPD1*, *PRAMEF,* and *LILR*) and all variants present in dbSNP except for those included in COSMIC somatic mutations (https://ftp.ncbi.nlm.nih.gov/snp/others/rs_COSMIC.vcf.gz) or the International Cancer Genome Consortium (ICGC) (https://ftp.ncbi.nlm.nih.gov/snp/others/snp_icgc.vcf.gz) variant lists. We used the Integrative Genomic Viewer (IGV) (83) to manually inspect a selection of MPNST related genes (*NF1*, *CDKN2A*, *SUZ12*, *EED*, *TP53*, *PTEN*, *RB1, NF2, SMARCB1, NRAS, BRAF, NTRK1, NTRK2* and *NTRK3*).

### Mutational Signatures

Since no true somatic calling was possible due to the lack of paired normal samples, we applied a series of filters to approximate a somatic call set: we filtered out the variants in dbSNP (except for those present in COSMIC or ICGC), variants with a population frequency (AF_popmax) higher than 1%, called in more than one individual, with a variant allele frequency (VAF) lower than 0.1, and variants in highly variable genes (*MUC3A*, *MUC5AC*, *OR52E5*, *OR52L1*, *SMPD1*, *PRAMEF*, and *LILR*). We used this call set enriched in somatic variants with the mutSignatures R package (84) to estimate the contribution of each of the COSMIC single base substitution (SBS) mutational signatures v3.3 to the mutational profile of each sample.

### Copy-Number and LOH analysis

We called copy-number alterations from mapped WGS data using CNVkit (85) with the recommended settings for WGS. We used a panel of normals for each dataset, added a region black-list (https://github.com/Boyle-Lab/Blacklist/blob/master/lists/hg38-blacklist.v2.bed.gz), the -no-edge option and 1000 bp bins for the CNV calling. Exact copy number profiles were called with the threshold method and we provided the Strelka2 germline results for the detection of LOH regions. To adjust the thresholds for tumor purity and ploidy, we first called CNVs assuming a 2n ploidy and 100% of tumor fraction. Then, based on these results and a pseudo-BAF obtained from the Strelka2 germline results using *loadSNPDataFromVCF* from CopyNumberPlots R package (86), we calculated the purity of each sample based BAF shift of copy-neutral LOH or heterozygous loss regions. With that, we considered a range of ploidies (from 2n to 4n) and manually selected the most accurate CNV calling. Finally, we annotated the CNV regions using biomaRt R package (87). Copy number alterations were plotted using the CopyNumberPlots (86) and karyoploteR (88) R packages.

### Structural Variant Calling

We used LUMPY (89) via Smoove (https://github.com/brentp/smoove) as the structural variant (SV) caller excluding the problematic regions defined in https://github.com/halllab/speedseq/blob/master/annotations/exclude.cnvnator_100bp.GRCh38.20170403.bed. For selected genes (*NF1*, *CDKN2A*, *SUZ12*, *EED*, *TP53*, *PTEN*, *RB1, NF2, SMARCB1, NRAS, BRAF, NTRK1, NTRK2* and *NTRK3*), we also performed a thorough visual inspection using the Integrative Genomic Viewer (IGV) (83) to detect additional breakpoints. To discard germline structural variants, we filtered out SVs present in two or more normal samples, present in the Database of Genomic Variants (DGV) (90), and the SVs present in two or more individuals. The representation of the structural variants was done using karyoploteR (88), CopyNumberPlots (86), and Circos (91).

SV type frequency was computed as the number of breakpoints associated to each SV type divided by the total number of SV breakpoints. An SV was considered to affect *CDKN2A* if at least one of its breakpoints overlapped *CDKN2A* locus +/- 1Mb. An SV was considered a hotspot SV if at least one of its breakpoints overlapped chr9:21971100-21972200. To compute the portion of genome affected by SVs, we flanked SV breakpoints by +/- 1Mb, merged the overlapping regions, and divided the sum of the width of these regions by the length of the genome.

### Fusion Genes

We used STAR-Fusion (92) for the detection of potential fusion genes from RNA-seq data. We intersected STAR-Fusion results with SV breakpoints and annotated them with the list of cancer fusion genes at https://cancer.sanger.ac.uk/census.

### Differential expression and feature selection

Differential gene expression was performed only with samples from the GeM consortium dataset to avoid batch effects. We used DESeq2 (93) to perform a pairwise comparison between G1-G4 groups and also G1G2 vs G3G4. We joined the results of these comparisons and used that set of genes as input for the Boruta (94) algorithm (maxRuns = 5000) to select relevant features. We then applied Recursive Feature Elimination with Cross-Validation (RFECV) over five independent iterations to reduce redundancy. Feature selection and performance analyses were conducted using the Boruta and pROC packages in R. We used clusterProfiler (10.18129/B9.bioc.clusterProfiler) to determine the enriched Biological Processes (BP) of Gene Ontology.

### Statistical Analysis

Box plots were created with R and statistical differences between groups were evaluated with t-test. Kaplan Meier curves were calculated using survival and ggsurvfit R packages. P-values < 0.05 were considered statistically significant.

## DATA AVAILABILITY

Data generated or used in this work has been deposited in the appropriate repositories and is publicly available. Data from the local cohort is available at the NF Data Portal (https://nf.synapse.org/) (*NOTE: Synapse accession number still in process of acceptance. Data has been deposited in the relevant repositories and concentrated in the NF Data Portal Project Page*). All data from the GeM consortium is available at EGA (EGAD00001008608).

## AUTHOR CONTRIBUTIONS

Conceptualization (MM-L, ES, BG); methodology and investigation (MM-L, JF-R, HM, IU-A, SO-B, EC-B, JF-C, AM, ER, MS, CR,RA, TS, IG, ES, BG); analysis (MM-L, JF-R, HM, IU-A, SO-B, EC-B, JF-C, AM, ER, MS, CR,RA, TS, IG, JCL-G, AC, AH-G, GT, MS, MCU, IB, CV, CR, HS, CL, MC, ES, BG); data curation (MM-L); visualization (MM-L, ES, BG); supervision (ES, BG); resources (JCL-G, AC, AH-G, GT, MS, MCU, IB, CV, CR, HS, CL, MC, ES, BG); funding acquisition (ES, BG); project administration (ES, BG); writing–original draft (MM-L, ES, BG); reviewing (all authors); writing–review and editing (MM-L, MC, ES, BG).

## ACKNOWLEDGMENTS

This work has been supported by the Instituto de Salud Carlos III National Health Institute - [PI20/00228; PI23/00583; PI23/00422] Plan Estatal de I+D+I 2013–2016, co-financed by the FEDER program – a way to build Europe; and Fundació La Marató de TV3 (51/C/2019). The work has also been partially supported by Fundación Proyecto Neurofibromatosis and the Generalitat of Catalonia and CERCA Program (2021 SGR 00967). MM-L is a YIA postdoctoral fellow of the Children’s Tumor Foundation (2025-01-004). We are indebted to Dr. David Miller and the GeM consortium for facilitating the access to the generated data (Cortes-Ciriano et al. 2023). We are also indebted to the “Biobanc de l’Hospital Infantil Sant Joan de Déu per a la investigació” integrated in the Spanish Biobank Network of ISCIII for the sample and data procurement. We thank the IGTP core facilities and their staff for their contribution and technical support. We are also grateful to the Fundación Proyecto Neurofibromatosis, the Asociación de Afectados de Neurofibromatosis (AANF), and the Catalan Neurofibromatosis Association (ACNefi) for their constant support.

