## Supplementary Figures for "Identification of Malignant Peripheral Nerve Sheath Tumor subtypes with distinct genomic identities"

**Supplementary Figure S4.** Copy-number profile of tumor samples without copy number adjustment by tumor purity and ploidy. Tumors are organized according to their percentage of tumoral cells. Warm colors represent gains (the darker the color, the higher the copy-number gain), and green colors represent losses (dark green represents homozygous loss). Loss of heterozygosity (LOH) is represented with a blue line under the copy-number regions. The top part of the graph represents the accumulation of losses, gains, and LOH regions in the different clusters. .......... 5
**Supplementary Figure S5.** Copy number profile statistics showing the differences between C1 and C2. **A**) Percentage of even copies and odd copies in the genome by cluster. **B**) Boxplot representing the percentage of Loss of Heterozygosity (LOH) present in the whole genome by cluster. **C**) Percentage of copy-neutral-LOH (CN-LOH) present in the whole genome by cluster. **D**) Percentage of the recurrent CN-LOH regions in CN-LOH. Statistical significance was considered when p-values < 0.05.E) Schematic representation of the genomic events leading to an MPNST formation. Image made using Biorender .......... 6
**Supplementary Figure S11.** Enriched biological processes (BP) terms of **A**) G1G2 vs G3G4 and **B**) G1 vs G2. 12

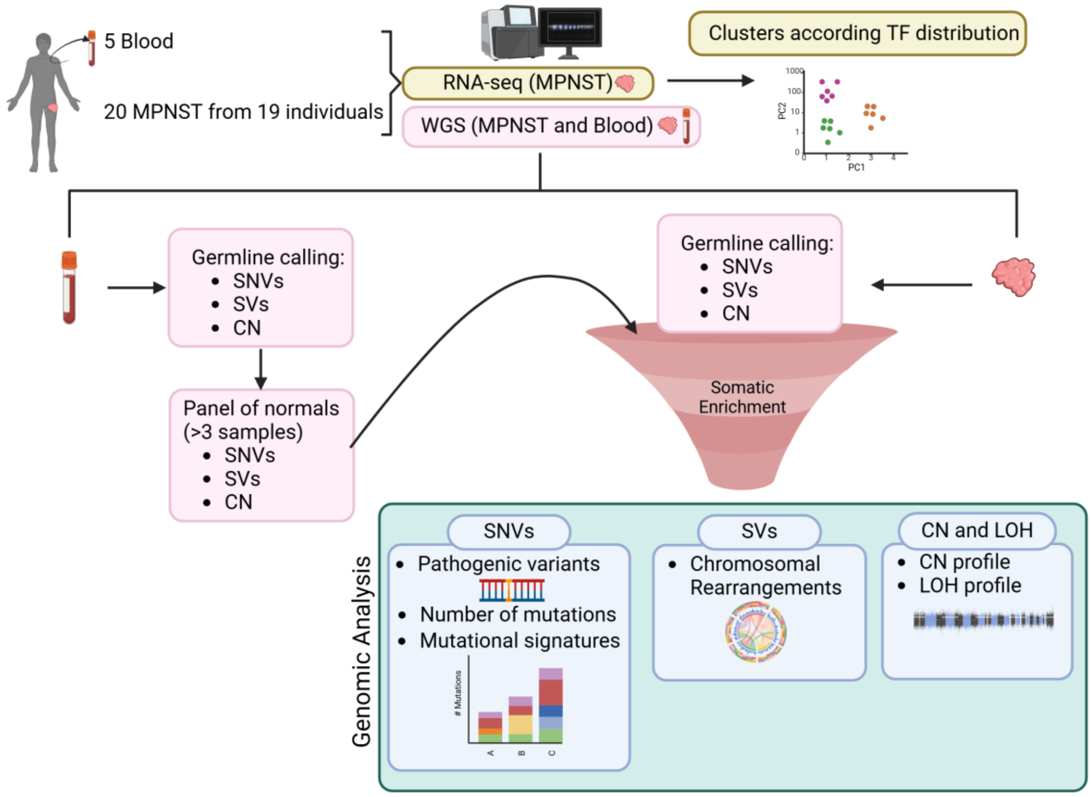

**Supplementary Figure S1.** Analysis pipeline applied to our sample set. Single-nucleotide variants (SNVs); Structural variants (SVs); Copy number (CN); Whole genome sequencing (WGS). Transcription factors (TF). Image made using Biorender

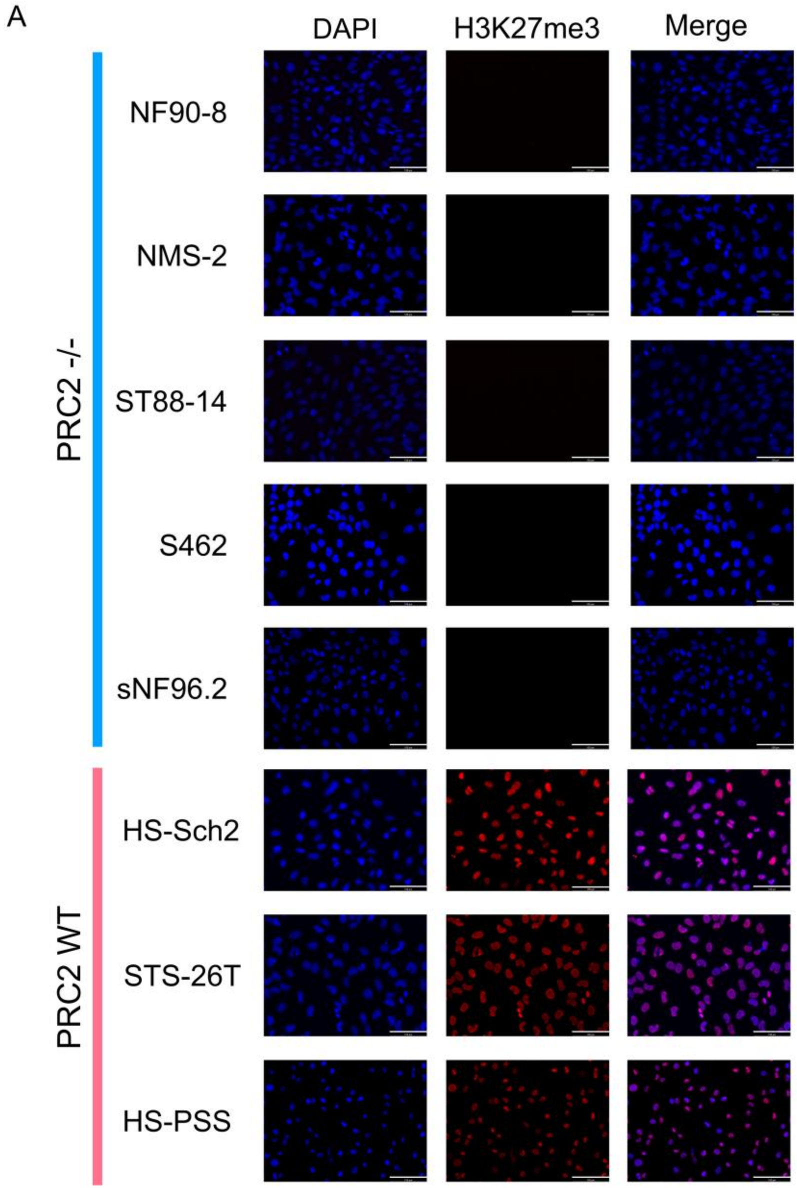

**Supplementary Figure S2.** Immunocytochemistry of H3K27me3 in five genuine MPNST cell lines with genomic inactivation of PRC2 (PRC2 -/-), and 3 cell lines with PRC2 wild type (WT). Scale bar: 100um

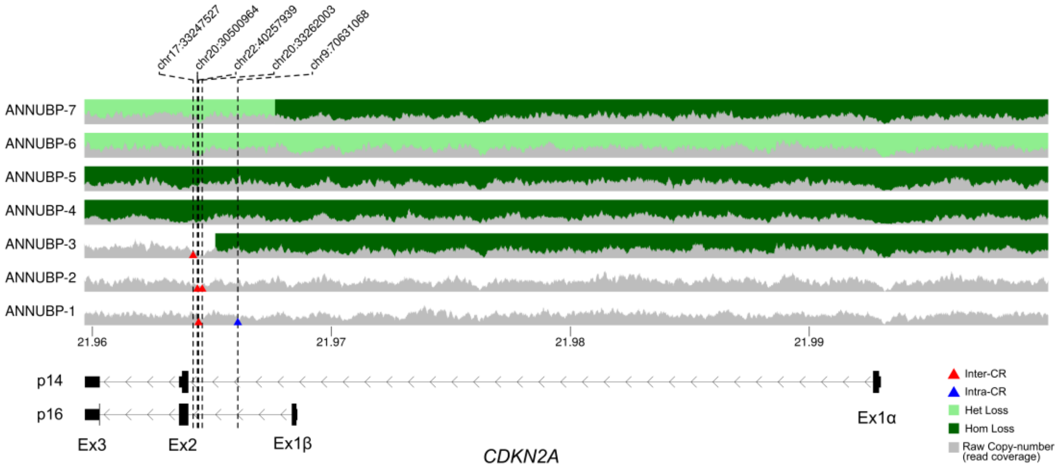

**Supplementary Figure S3.** Inactivation of *CDKN2A* in ANNUBPs. Breakpoints inactivating *CDKN2A* are depicted with triangles.

A

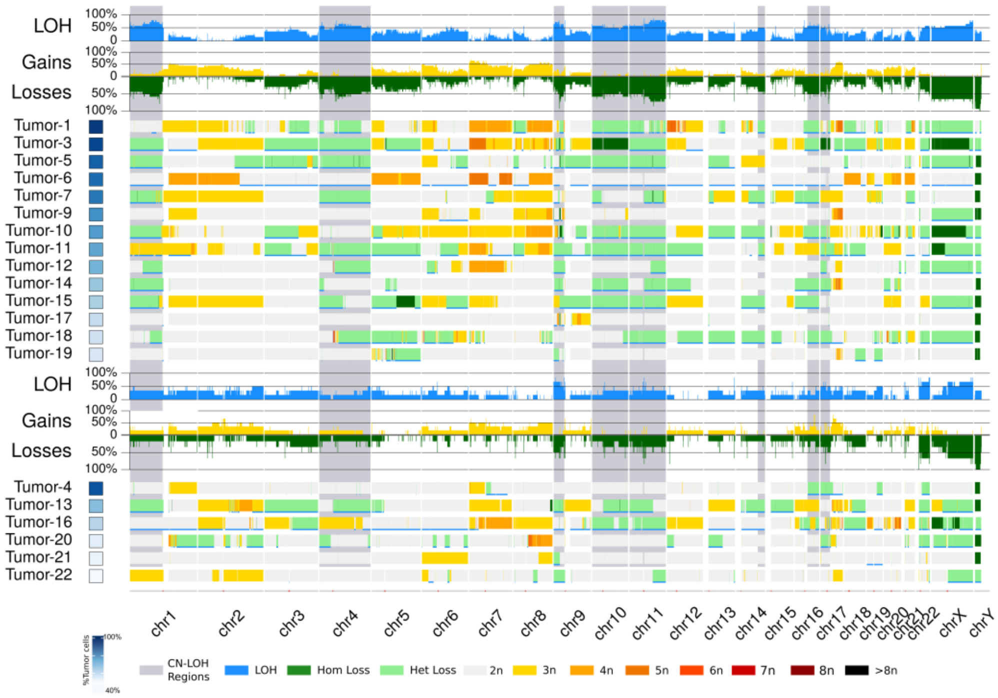

**Supplementary Figure S4.** Copy-number profile of MPNST samples without calling adjustement based on purity and ploidy. Tumors are organized in two groups (top C1, bottom C2) and inside each group according to their percentage of tumoral cells. Warm colors represent gains (the darker the color, the higher the copy-number gain), and green colors represent losses (light green heterozygous, dark green homozygous). Loss of heterozygosity (LOH) is represented with a blue line under the copy-number . The top part of each group represents the accumulation of losses, gains, and LOH.

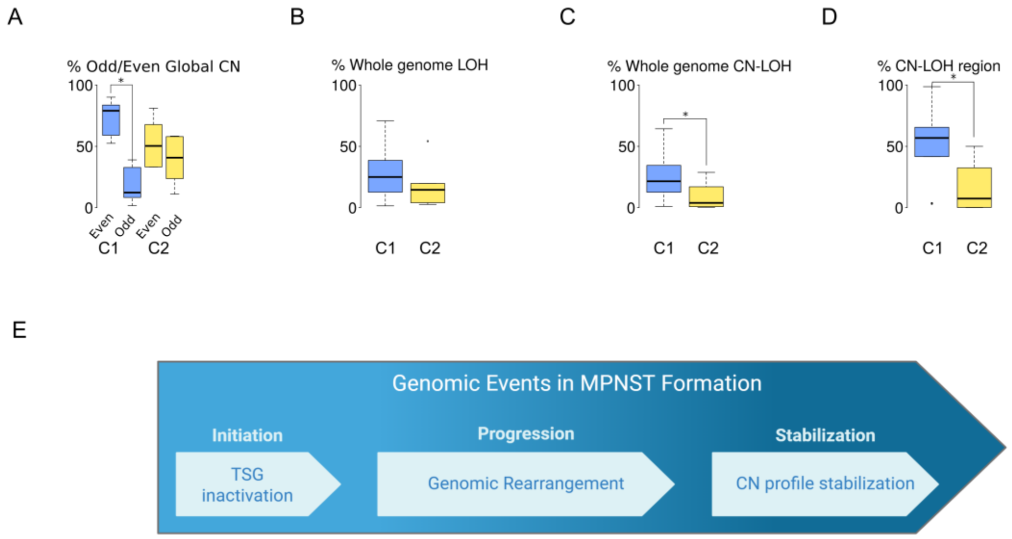

**Supplementary Figure S5.** Copy-number and Loss of Heterozygosity (LOH) differences between C1 and C2. **A**) Percentage of the genome at odd or even copy-number state. **B**) Boxplot representing the percentage of genome with LOH. **C**) Percentage of genome with copy-neutral-LOH (CN-LOH). **D**) Percentage of the regions identified as recurrent CN-LOH actually in CN-LOH. Statistical significance was considered when p-value $< 0.05$. **E)** MPNST formation model with three steps, Initiation, Progression and Stabilization. Image made using Biorender

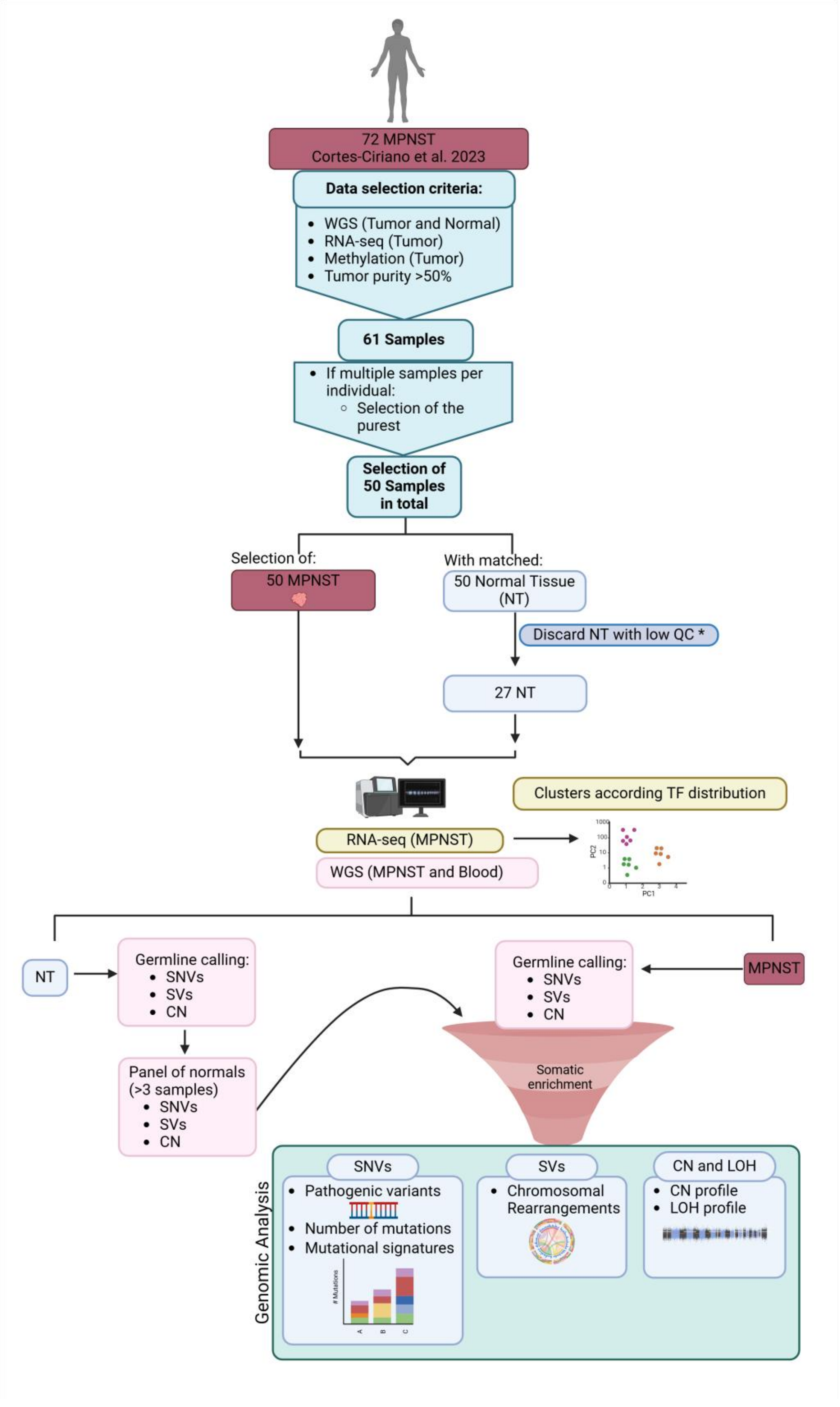

**Supplementary Figure S6.** Sample selection criteria and analysis schema of GeM Consortium samples (Cortes-Ciriano et al. 2023). Single-nucleotide variants (SNVs); Structural variants (SVs); Copy number (CN); Whole genome sequencing (WGS); Transcription factors (TF). * low QC examples in Supplementary Figure 7. Image made using Biorender

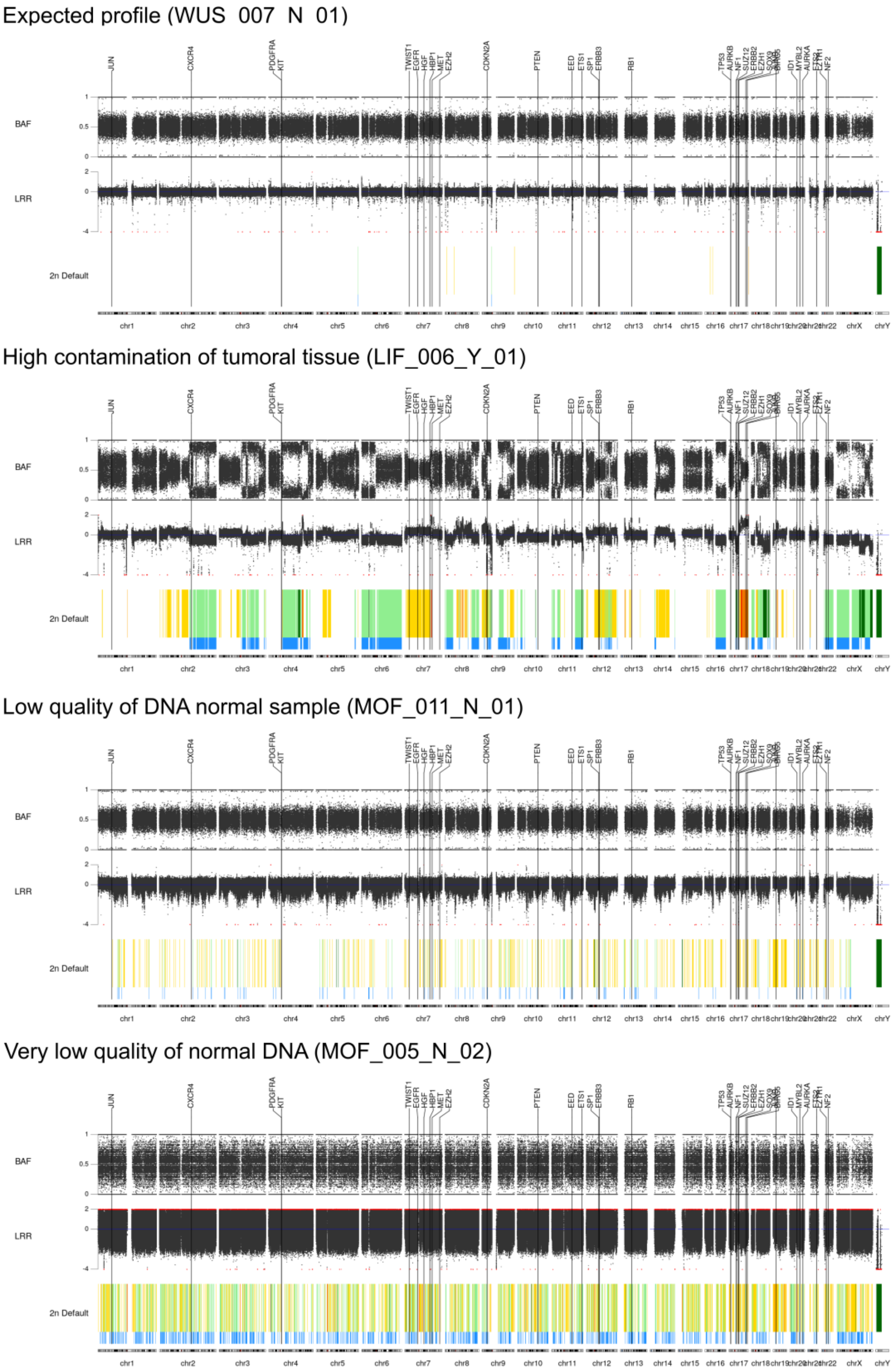

**Supplementary Figure S7.** Examples of low-quality normal samples found in the GeM consortium set (Cortes-Ciriano et al., 2023).

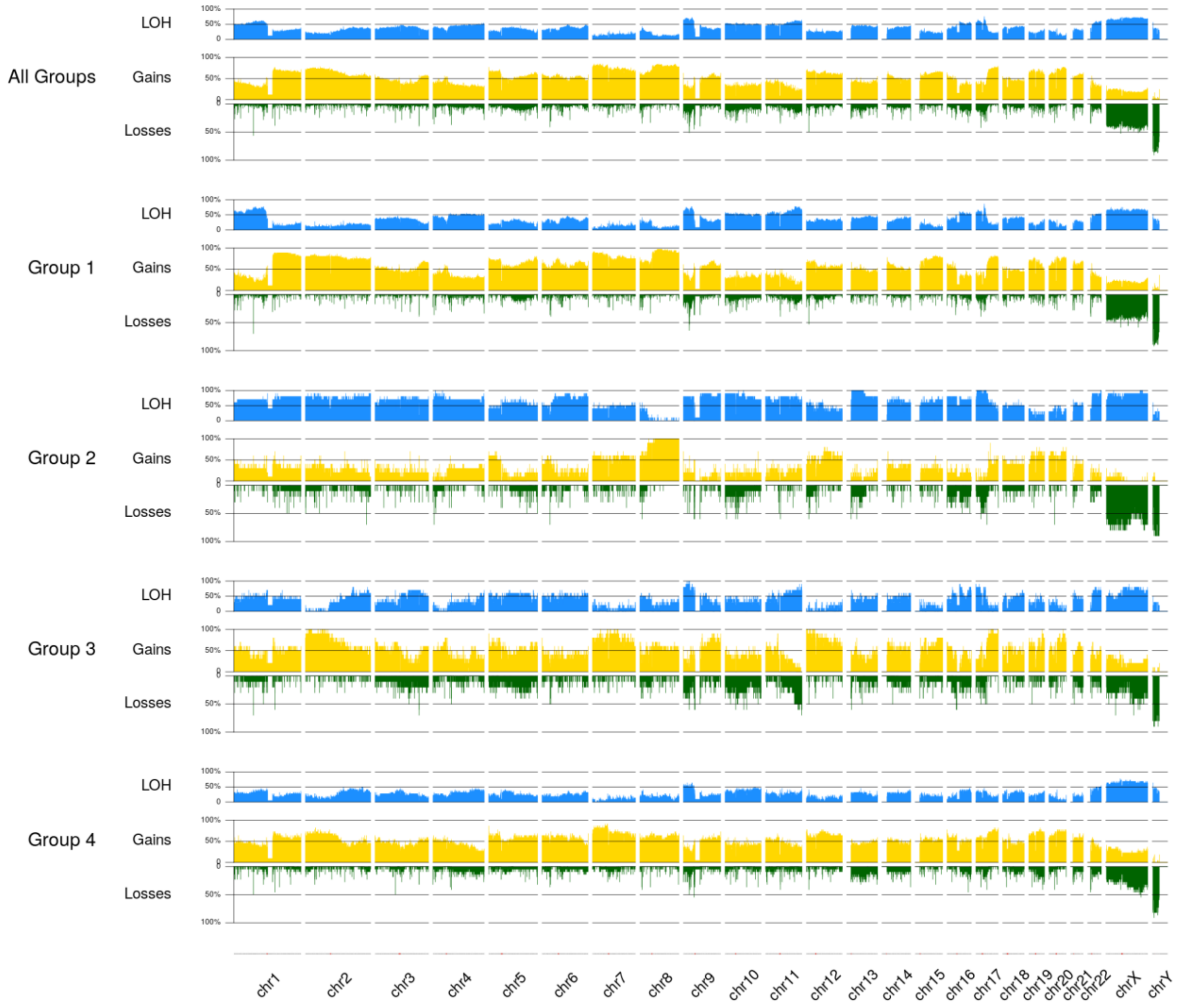

**Supplementary Figure S8.** Summary of the copy-number profile of the different groups. For each group, the proportion of tumors with loss of heterozygosity (LOH) is represented in blue, gains in yellow and losses in green.

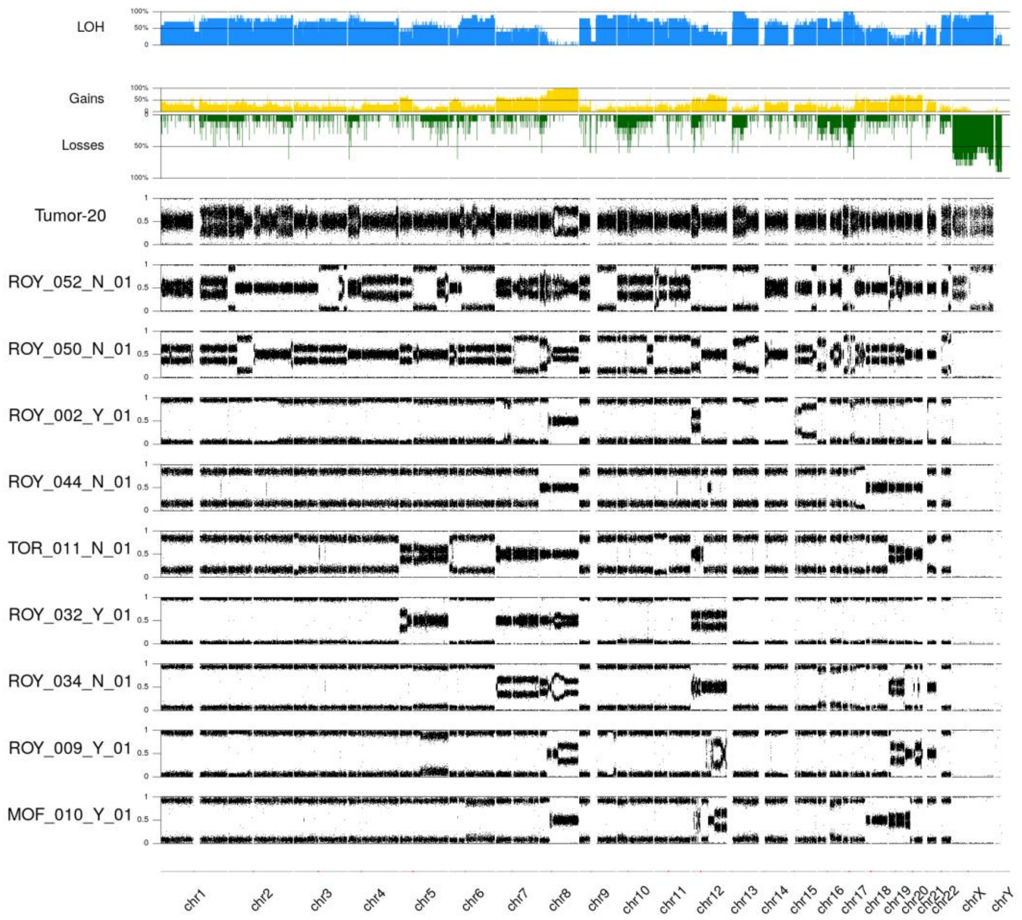

**Supplementary Figure S9.** B-allele frequency (BAF)-like profile of the MPNSTs in G2. the proportion of tumors with loss of heterozygosity (LOH) is represented in blue, gains in yellow and losses in green..

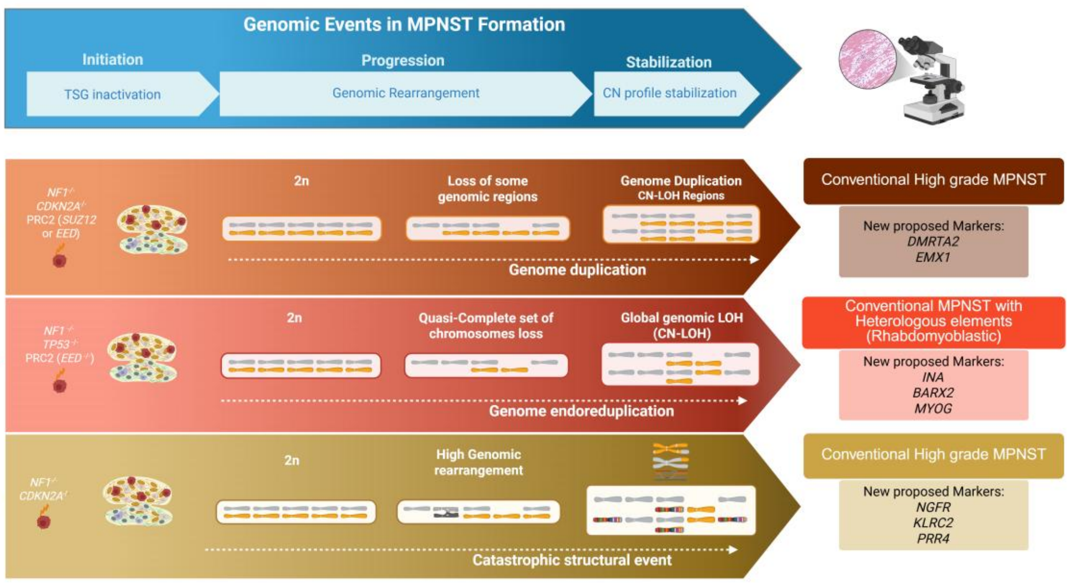

**Supplementary Figure S10.** Schematic representation of the distinct MPNST formation paths of G1, G2 and G3, including the specific set of TSG inactivated and the genomic rearrangements driving the progression and stabilization. It also includes the most frequent histological classification of tumors in each group and the proposed gene markers for differential diagnosis. Image made using Biorender

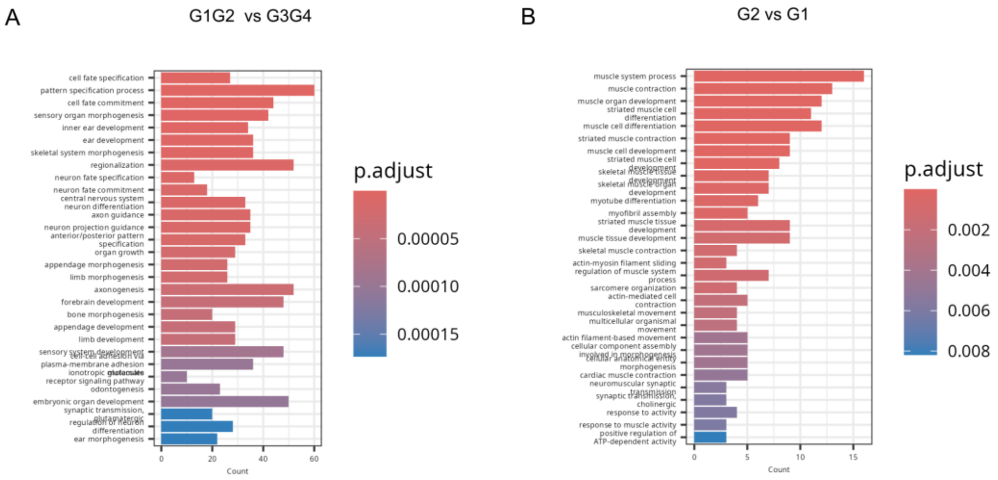

**Supplementary Figure S11.** Enriched biological processes (BP) of **A**) G1-G2 vs G3-Others and **B**) G1 vs G2.
