## Supplementary File 1 for "Identification of Malignant Peripheral Nerve Sheath Tumor subtypes with distinct genomic identities"

### Supplementary File 2

The “Others” group seems to contain different tumoral entities. In this file we provide several pieces of evidence of why we considered that some tumors are other entities instead of MPNSTs. Tumor-21 and Tumor-22 were in the “Others” group, and as we already mentioned, they seem to be tumoral entities associated with melanoma (**Manuscript Figure 2**) due to the presence of activating mutations and also a high number of mutations with SBS7 signature associated.

Some examples of potentially non-MPNST tumors:

- **ROY_047_N_01**, in which we detected *FUS-TFCP2* fusion gene already described in Rhabdomyosarcomas (Fig. 1A), and also high *MYOD1* expression (Fig. 1B), which is also associated with this type of tumor (1), and presented Rhabdomyoblastic (triton) traits (**Figure 1**). These pieces of evidence lead us to think it might had been misidentified at the moment of the diagnosis.

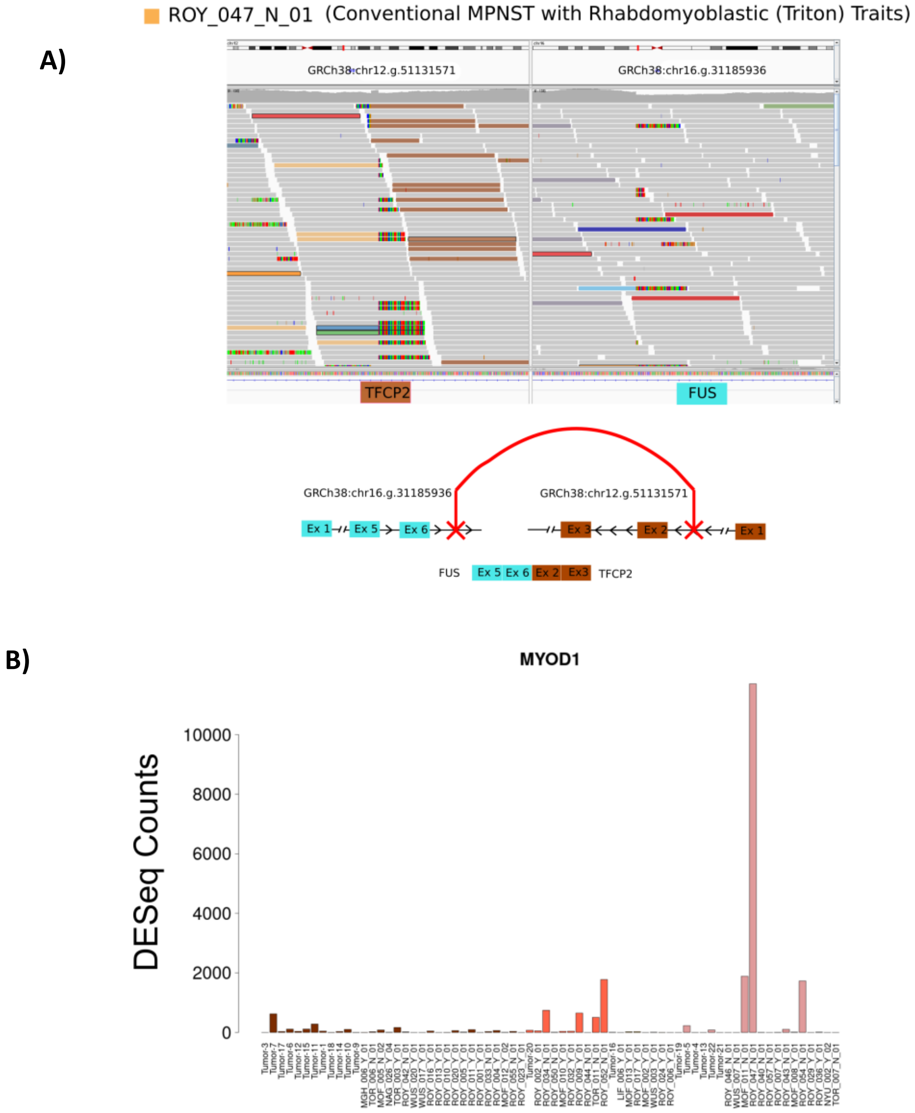


**Figure 1**. Pieces of evidence associated with ROY_047_N_01 tumor.

- **ROY_043_N_01** bore another fusion gene (*ZFP64-NCOA3*) also described in a set of spindle cell rhabdomyosarcoma (2) (**Figure 2**). As described by Han and colleagues, the expression of *MYOD1* was also weak in this sample (**Figure 1**). All these pieces of evidence and the presence of rhabdomyoblastic histology led us to think it was misidentified at the moment of the diagnosis.

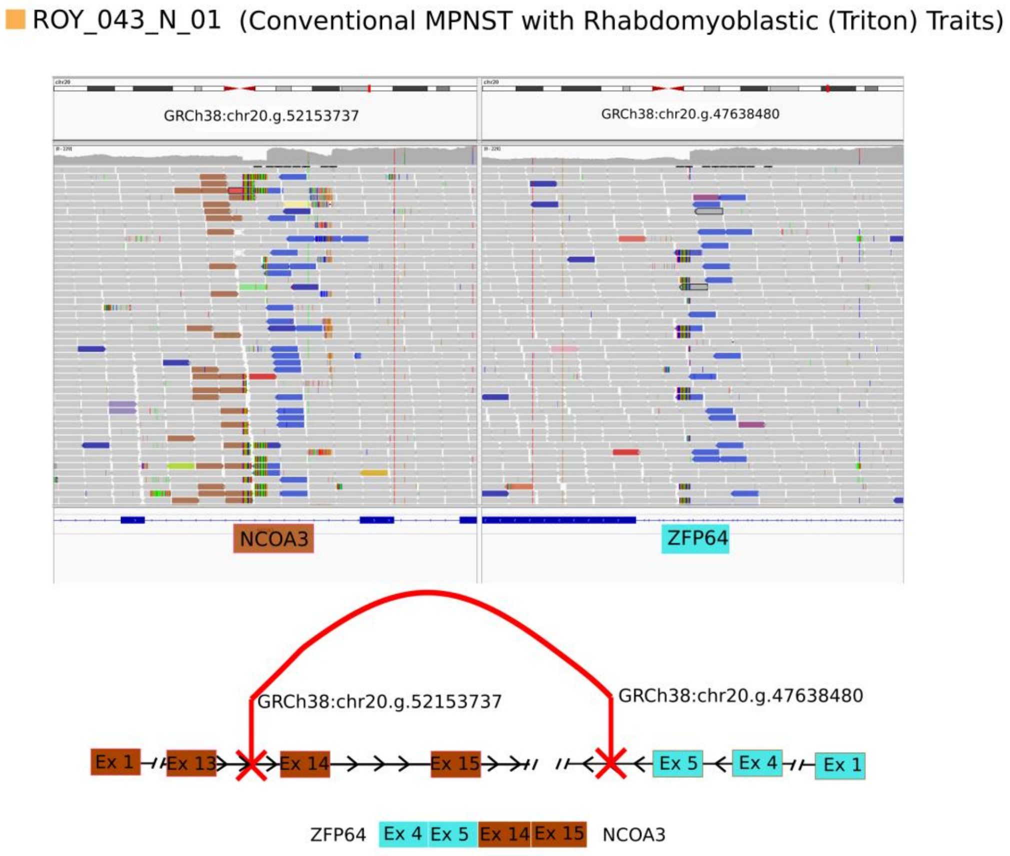


**Figure 2.** Pieces of evidence associated with ROY_043_N_01 tumor.

Finally, we also detected some NF1 samples with classic MPNST histology but bearing *NTRK* fusion genes (ROY_036_Y_01 and Tumor-13).

- **ROY_036_Y_01** bore *AKAP13-NTRK3* fusion gene and presented remarkably gained expression of *NTRK3* compared with the rest of the samples.

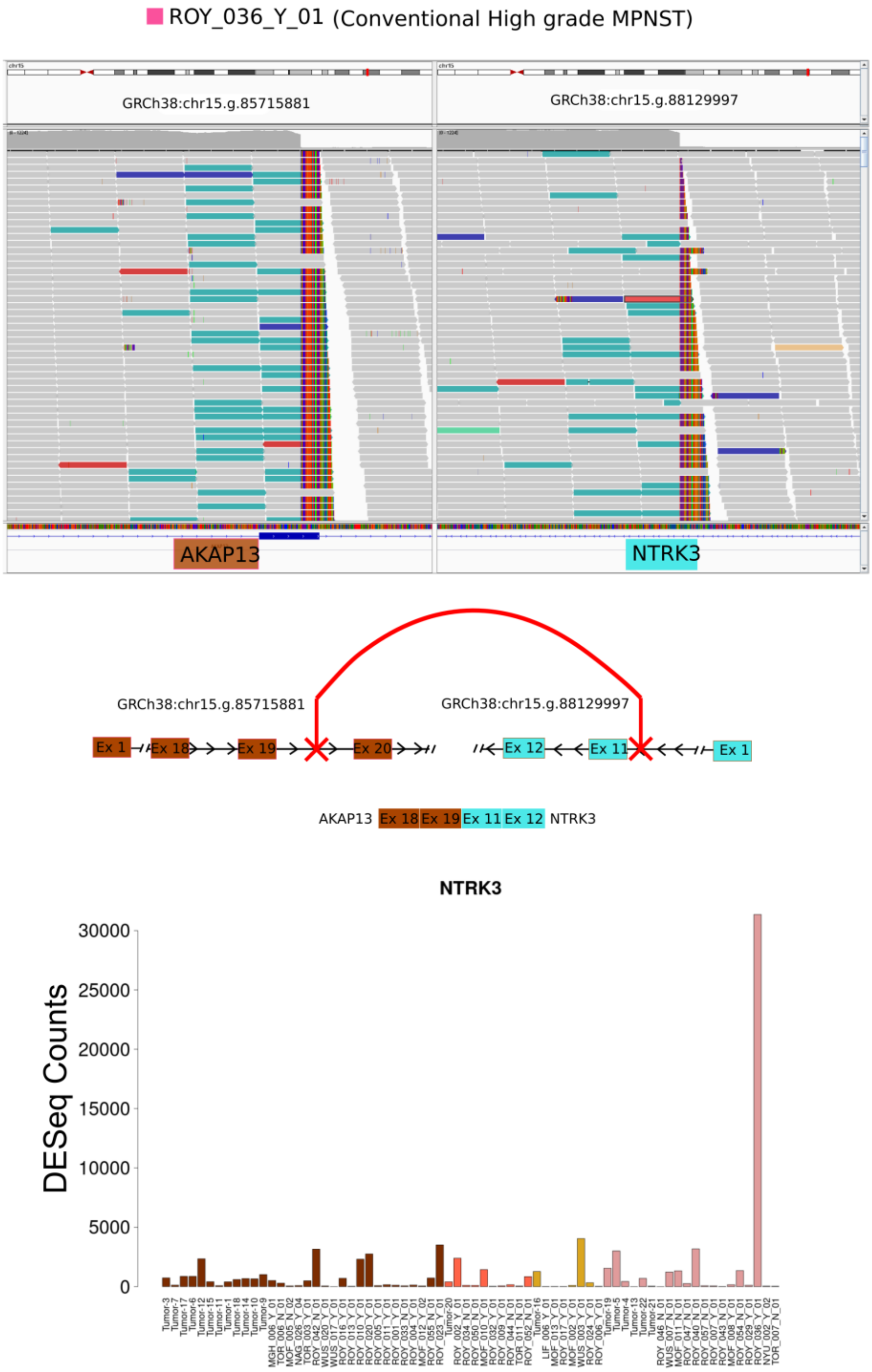


**Figure 3.** Pieces of evidence associated with ROY_036_N_01 tumor.

- **Tumor-13** *LMNA-NTRK1* fusion gene was already described in Creus-Bachiller et al., 2024 (3), here named as SP-05. We detected a remarkable gain in NTRK1 expression compared with the rest of the samples (**Figure 4**).

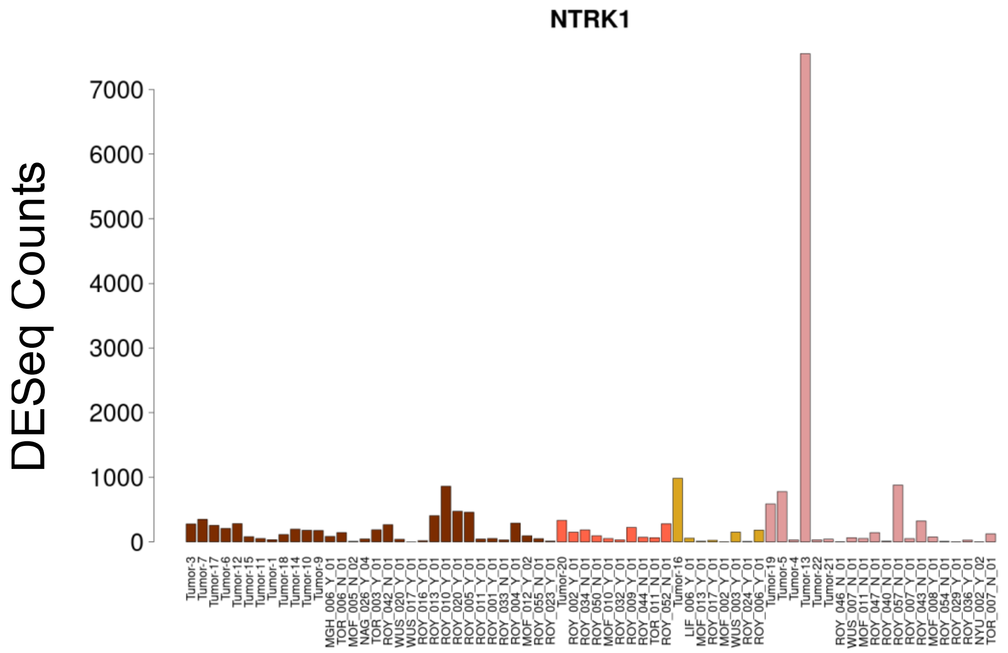


**Figure 4.** Pieces of evidence associated with Tumor-13 tumor.
